# Age-related inflammatory changes and perineuronal net dynamics: implications for neurodegenerative disorders

**DOI:** 10.1101/2025.09.10.675190

**Authors:** Zachary A. Colon, Shannon C. Chan, Kathleen A. Maguire-Zeiss

## Abstract

**Background:** Healthy aging alone can lead to cognitive decline, decreased brain size, protein aggregation, accumulation of senescent cells and neuroinflammation. Furthermore, age is the primary risk factor for several neurodegenerative disorders such as Parkinson’s and Alzheimer’s disease. Age-related neuroinflammation, as known as inflammaging, is thought to restrict brain plasticity. Perineuronal nets (PNNs), specialized extracellular matrix structures surrounding fast-spiking parvalbumin (PV) interneurons, regulate plasticity and protect neurons from oxidative stress. Given the known impact of inflammaging on neural circuits, this study examines age-associated changes in PNN homeostasis, glial activation, and neuroinflammation in two brain regions relevant to age-related neurodegenerative diseases.

**Methods:** We analyzed young (4-month-old) and aged (22-month-old) C57BL/6J male mice for several behavioral phenotypes [hippocampal-dependent spatial learning using the Barnes maze; locomotion and anxiety-related behaviors using Open field and T-maze]. Using immunostaining, PNNs (*Wisteria floribunda* agglutinin and aggrecan), PV interneurons, and microglial activation (Iba1) were quantified in both the hippocampus and dorsal striatum. Glial morphology was examined using a battery of cell body, branching, and endpoint analyses. Quantitative RT-PCR was used to analyze changes in the gene expression of inflammatory and extracellular matrix markers.

**Results:** Aged mice exhibited hippocampal-dependent memory deficits without alterations in locomotion or anxiety-related behavior. PNN counts increased in the aged hippocampus, particularly in CA2, with a higher proportion of WFA^+^ and aggrecan^+^ PNNs. In contrast, PNN homeostasis was maintained in the dorsal striatum. In general, Aged mice showed increases in microglial activation and a subset of inflammatory markers. We report brain region- and age-specific gene expression changes in complement, matrix metalloproteinases, and other inflammatory markers. Aged striatal microglia displayed an activated morphology with larger cell bodies and reduced branching, as well as increased expression of markers for microgliosis (Iba1, TREM2, CD68).

**Conclusions:** These findings suggest that aging differentially affects neuroinflammation and PNN integrity across brain regions. The hippocampus exhibits PNN accumulation, neuroinflammation, and behavioral changes, whereas the striatum maintains PNN homeostasis concurrent with increased microglial activation. This work suggests that neuroinflammation contributes to age-related changes in PNNs and behavior underscoring the importance of region-specific therapeutic strategies targeting PNN regulation.

## Background

Age remains the predominant risk factor for several neurodegenerative disorders (1,2). Despite this, several models of neurodegenerative disorders fail to incorporate the effects of normal aging on the brain. According to the World Health Organization, the global population of individuals aged 60 and older is projected to double to 2.1 billion by 2050 (3), presenting age as an expected risk factor to grow in the upcoming years. This demographic shift underscores the urgent need to understand the effects of normal brain aging.

Inflammaging is the chronic, low-grade inflammation that typically accompanies aging (4). The term "inflammaging," a combination of "inflammation" and "aging," describes the systemic inflammatory state observed in elderly individuals that is thought to also contribute to age-related disease pathology (4). This state is characterized by elevated levels of pro-inflammatory cytokines and sustained activation of the innate immune system. Inflammaging represents a significant factor contributing to the pathogenesis of age-related diseases, including neurodegenerative disorders, cardiovascular diseases, and metabolic syndrome, although it is unclear if these diseases are a manifestation of inflammaging or simply an acceleration in age-related disease pathology (4). Recent human plasma proteomics analyses have identified age-related alterations to the brain and immune system as significant indicators of mortality and longevity (5). Several mechanisms contribute to inflammaging. For example, senescent cells accumulate with age and secrete pro-inflammatory cytokines, chemokines, and proteases, known as the senescence-associated secretory phenotype (2). Immunosenescence causes the aging immune system to undergo functional changes, including reduced adaptive immune responses and increased activation of the innate immune system, resulting in a heightened inflammatory state. In addition, age-related organelle dysfunction leads to increased production of reactive oxygen species (ROS), protein accumulation and activation of inflammatory signaling pathways (6,7). Likewise, persistent infections or reactivation of latent pathogens will drive chronic immune activation and inflammation. Whether intrinsic or extrinsic, these factors all contribute via multiple mechanisms to an age-related neuroinflammatory profile in the brain.

As chronic inflammation is one hallmark of aging (reviewed in (7)) it is not surprising that circulating levels of cytokines, like IL-6, have been identified as biomarkers for inflammaging associated with an increase in mortality (8). In terms of the brain, less is known about the neuroinflammaging profile. In rodent models, studies show that inhibition of TNF-alpha signaling, the inflammasome, and prostaglandin signaling all improve cognition, motor performance, and the neuroinflammatory milieu (9–11). This along with the fact that aging is the greatest risk factor for two common age-related neurodegenerative disorders, Parkinson’s and Alzheimer’s disease, drove us to investigate the neuroimmune profile of brain regions associated with these disorders (12).

Interestingly, one common biological process associated with immune system aging, brain aging, and brain function is extracellular matrix regulation (ECM; (13,14)). Failure of the brain to properly maintain ECM modulation results in a decrease in brain circuit plasticity (reviewed in (15)). It was recently reported, in a large human plasma proteomic study, that changes in circulating extracellular matrix protein levels are associated with brain aging (5), in particular this study found an association with increased brevican and decreased matrix metalloproteinase-9 (MMP9) in ‘youthful brain agers’. Brevican is found in perineuronal nets (PNNs), a specialized extracellular matrix structure which enwraps the cell body and proximal dendrites of neurons providing structural support, regulating plasticity, and protecting the neuron from oxidative stress (16–18). PNNs are sculpted by a class of endopeptidases called MMPs which degrade various components of the matrix (19) and can be upregulated in neuroinflammation. Furthermore, PNN components and MMPs are made by glia and altered in various disease states.

PNNs often envelop fast-spiking parvalbumin (PV) interneurons and these neurons are present in both the striatum and hippocampus. These interneurons are important for maintaining the excitatory/inhibitory balance required for the timing and synchronization of neuronal circuits through modulation, sparse encoding, and pattern separation (20). Dysfunction of these cells has previously been linked to epilepsy, schizophrenia, and autism (21,22), as well as PD (23) and AD (24). Furthermore, both aging and neuroinflammation are implicated in PNN dysfunction and changes in PNNs can alter the synaptic plasticity of PV interneurons (25), so the homeostasis of PNNs is hypothesized to be important for the maintenance of neuronal circuits.

Little is known regarding the combination of regional differences in PNNs, neuroinflammation, and PV neuron numbers in aging. Therefore, in this study we utilized 4-month-old adult mice (Young) and 22-month-old mice (Aged), an age equivalent to humans aged 60-70 years (26), to investigate age-associated inflammation, PNN alterations, and PV neuron number in the dorsal striatum and hippocampus. Here we report both age-related and region-specific changes in neuroinflammation, PNN homeostasis, and glial morphology. Taken together, this work suggests that aging has differential effects in the brain, with some areas more vulnerable to inflammaging than others.

## Material and Methods

### Animals

C57BL/6J male mice (Jackson Laboratories, JAX#000664) were housed in the Georgetown University Division of Comparative Medicine facilities under a 12 h light/dark cycle with food and water available ad libitum. Cohorts of mice were aged to 4-months (Young) or 22-months (Aged). Five animals per group (N=5) were used in accordance with the Georgetown University Institutional Animal Care and Use Committee and National Institutes of Health ethical guidelines.

### Behavioral Testing

Behavioral testing was conducted in the Georgetown University Department of Neuroscience behavioral facilities between 9:00 AM and 5:00 PM. Mice were habituated to handling and testing locations for one week before behavioral assessments. Testing occurred over 14 days before tissue collection, with a randomized testing order.

### Barnes Maze

Spatial learning and memory were assessed using a standard Barnes maze (92 cm diameter, 20 holes, Stoelting Co.) with four fixed extra-maze visual cues (triangle, square, circle, star). Mice were trained over 4 consecutive days with 3 trials per day (5 minutes/trial, 60-minute inter-trial intervals). If a mouse failed to locate the escape hole, it was guided into it at the end of the trial and allowed to remain there for 30 seconds. Twenty-four hours after the final training session, a 5-minute probe trial was conducted with the escape box removed. Behavior was recorded and analyzed for latency to the target hole, time spent in the target zone and each quadrant, and search strategy. Search strategies were characterized as the following: random (animal moves back and forth across the platform until the escape hole was found, serial (animals traveled around the periphery of the maze until they found the correct hole in a systematic pattern), or direct (animals moved to escape hole directly with less than two errors (27).

### Open Field

The Open field test assessed general activity, anxiety, and exploratory behavior. Mice were placed in a 45 cm × 45 cm × 50 cm enclosed arena for 5 minutes, and total distance traveled, speed, time spent in the inner zone versus the periphery, and rearing/grooming frequency were recorded. Increased inner zone activity indicated reduced anxiety.

### T-Maze

The T-maze was used to assess working memory and decision-making. Mice were placed in the start arm facing away from the decision point. Following a 180° turn, the start door was removed to begin the trial. Mice were allowed to freely choose one of the two goal arms. Upon entry, the selected arm was closed off, and the mouse remained in the arm for 60 seconds before being removed. The maze was cleaned with 70% ethanol between runs. Each trial consisted of two consecutive runs separated by a brief interval. An alternation was defined as the mouse entering the opposite arm in the second run relative to its initial choice. Failure to switch arms was recorded as a non-alternation. Each mouse completed three trials, and the number of alternations and decision times were recorded.

### Tissue Collection

Mice were anesthetized before intracardial perfusion with artificial cerebrospinal fluid (125 mM NaCl, 25 mM NaHCO_3_, 3.5 mM KCl, 1.25 mM NaH_2_PO_4_, 10 mM glucose, 1 mM MgCl_2_, and 2 mM CaCl_2_). Brains were hemisected, with one half fixed in 4%paraformaldehyde/4%sucrose (w/v) in phosphate buffer (pH = 7.2-7.4) for 24 hours before sucrose gradient saturation (20% (w/v) and 30% (w/v) sucrose in phosphate buffer). Tissue was sectioned (30-μm thick) using a sliding microtome (Model# HM-400R, MICROM GmbH) cooled using dry ice and 100% isopropanol to approximately -70°C to -78.5°C and stored in cryoprotectant at -20°C. The other half of the brain was microdissected, with the dorsal striatum and hippocampus flash-frozen at -80°C for RNA analysis.

### Quantitative Real-Time PCR

Total RNA was extracted from the dorsal striatum and hippocampus using the RNeasy Mini Kit (Qiagen, Catalog #74104) and transcribed into cDNA using the High-Capacity RNA-to-cDNA Kit (Applied Biosystems, Catalog #4368814). Gene expression of inflammatory, glial, and perineuronal net markers were quantified using TaqMan primer/probe pairs. 18S and GAPDH were used as reference genes. Fold changes were calculated using the ΔΔCt method, with statistical analyses performed on ΔCt values.

### Immunostaining

Free-floating brain sections were stained for perineuronal nets (PNNs), parvalbumin (PV) neurons, and glial cells using established protocols (32). PNNs were stained with Wisteria floribunda agglutinin (WFA)-fluorescein (Vector Labs, Catalog #FL-1351) or biotinylated-WFA (Vector Labs, Catalog #B-1355) depending on the experiment. Aggrecan was detected using anti-aggrecan (Millipore, Catalog #AB1031). PV neurons were labeled with an anti-parvalbumin antibody (Sigma-Aldrich, Catalog #P3088). Microglia were labeled with an anti-Iba1 antibody (FUJIFILM Wako, Catalog #019-19741). Secondary antibodies included Alexa Fluor-488 α-Rabbit (Invitrogen, Catalog #A-11008), Alexa Fluor-594 α-Mouse (Invitrogen, Catalog #A-21125), Alexa Fluor-594 α-Goat (Invitrogen, Catalog #A-11058), and Streptavidin-Cy5 (Invitrogen, Catalog #SA1011).

### Imaging and Analysis

Researchers were blinded to conditions for imaging and analysis. Exposure times and laser intensities were optimized and maintained across samples. Imaging for cell counts were performed using a Zeiss Axio Imager.Z2 microscope (20X objective). Images were stitched and processed using ZEISS ZEN (Blue Edition) software (Carl Zeiss Microscopy) (28) and FIJI (ImageJ) (29). Cell counts were performed using tiled stitched images of the regions of interest with 30% overlap and stitched via Zen Blue “Stitching” function. Regions of interest (ROI) were identified and outlined with the Allen Brain coronal mouse Atlas as a reference (30). Mean intensity was measured via FIJI’s ‘measure’ function. Cell counts were manually counted and averaged over the total area of interest. Single-cell analyses involved 15-μm z-stacks (0.24-μm steps) of PV and PNN colocalized cells.

### Single-cell Microglia Analysis

Microglia were imaged at 63X (15-μm stacks, 0.3-μm steps). Confocal z-stacks were converted to 8-bit “average intensity” 2D projections. Iba1 immunofluorescence (IF) intensity was quantified using the ‘Measure’ function for both whole images and individually outlined cell bodies, with mean intensity values calculated per region and per cell. For morphological analysis, images were background subtracted (rolling ball radius: 50 pixels), thresholded (97–97.5%), binarized, and processed with a fast Fourier transform (FFT) bandpass filter (50 pixels) and unsharp mask (radius: 1.0 pixel). Noise was reduced by despeckling and outlier removal, following protocols adapted from (31,32). Cell bodies were manually traced using the freehand selection tool on thresholded images, and area and perimeter measurements were obtained. To assess microglial process complexity, individual cells were isolated by cropping and clearing background, with fragmented processes manually reconnected. Sholl analysis was performed using the Sholl Analysis plugin (v4.2.1) (33) in FIJI (29) on binarized cells with the following parameters: starting radius = 0 μm, ending radius = 100 μm, step size = 5 μm, and five samples per radius. The number of intersections at each radius was recorded, and averages were calculated per region and animal. Endpoint analysis was conducted by counting the number of terminal points per cell to quantify branching complexity. Additionally, cells were characterized as either “highly branched” (>2 branches, ramified) or “minimally branched” (≤2 branches, amoeboid-like, bipolar, or rod-like) based on the number of observable projections. All quantitative measures, including IF intensity, soma morphology, Sholl intersections, and endpoints, were averaged across cells for each region and animal and used for group-level statistical comparisons.

### Colocalization Analysis

Colocalization was performed using the Just Another Colocalization Plugin (JaCoP) (34) in FIJI. PV^+^:WFA^+^, PV^+^:ACAN^+^, and WFA^+^:ACAN^+^ colocalization were quantified to assess interactions between parvalbumin neurons and perineuronal net components. Cells were randomly selected from prior cell counts and imaged at 40× magnification as 15-μm z-stacks with 0.24-μm step sizes. Images were preprocessed with background subtraction (rolling ball radius = 20 pixels) and “average intensity” 2D projections were created for each image. ROIs were drawn to isolate PV^+^ cells, WFA^+^-labeled PNNs, or ACAN^+^ PNNs. For each comparison, the selected ROI channel was loaded as Image A, and the colocalized target channel as Image B in JaCoP. Pearson’s correlation coefficient, Manders’ coefficients (M1, M2), and overlap coefficients (k1, k2) were calculated. Costes’s automatic thresholding was applied to minimize bias and ensure consistent thresholding across samples and Pearson’s correlation coefficient were calculated.

### Regression Analysis

The relationship between [PV], [WFA], and [ACAN] was assessed by Pearson’s correlation coefficient to examine the strength of association between the targets. Simple linear regression was performed using ordinary least squares fitting and statistical significance was determined by testing whether the slope of the regression differed from zero and if the slopes between groups were significantly different. For regression analyses, total intensity of regions of interests were calculated using the tiled stitched images and normalized to acquire an intensity value per animal for [WFA], [PV], and [ACAN]. Normalized animal intensity values were plotted on x-y coordinates with the independent variable [WFA] intensity along the x-axis and dependent variable intensities along the y-axis for each region of interest. Linear regression analysis was performed for each age group to calculate the coefficient of determination (R^2^) and a t-test for slopes of each region was performed to identify significant relationships between the variables.

### Statistical and Data Analyses

Normality of data distributions were assessed using GraphPad Prism’s "Normality and Lognormality Tests" module. The D’Agostino–Pearson and Anderson–Darling tests were applied, with the significance threshold set at α = 0.05. Unpaired Student t-tests, linear regression, coefficient differences t-test for slopes, one-way, two-way, or three-way ANOVAs with Tukey corrections were used where appropriate. Data are reported as mean ± SEM, with significance set at p < 0.05. Analyses were conducted using GraphPad Prism 10.2.3 (GraphPad Software, Boston, MA, USA). Complete data for **Figs. 1-4** and **Fig. 6** is available in **Supplementary Table 1,** with **Fig. 5** data available in **Supplementary Table 2**. qPCR primer sequences and catalog numbers can be found in **Supplementary Table 3.** Imaging resolution and metadata can be found in **Supplementary Table 4.**

**Figure 1.**
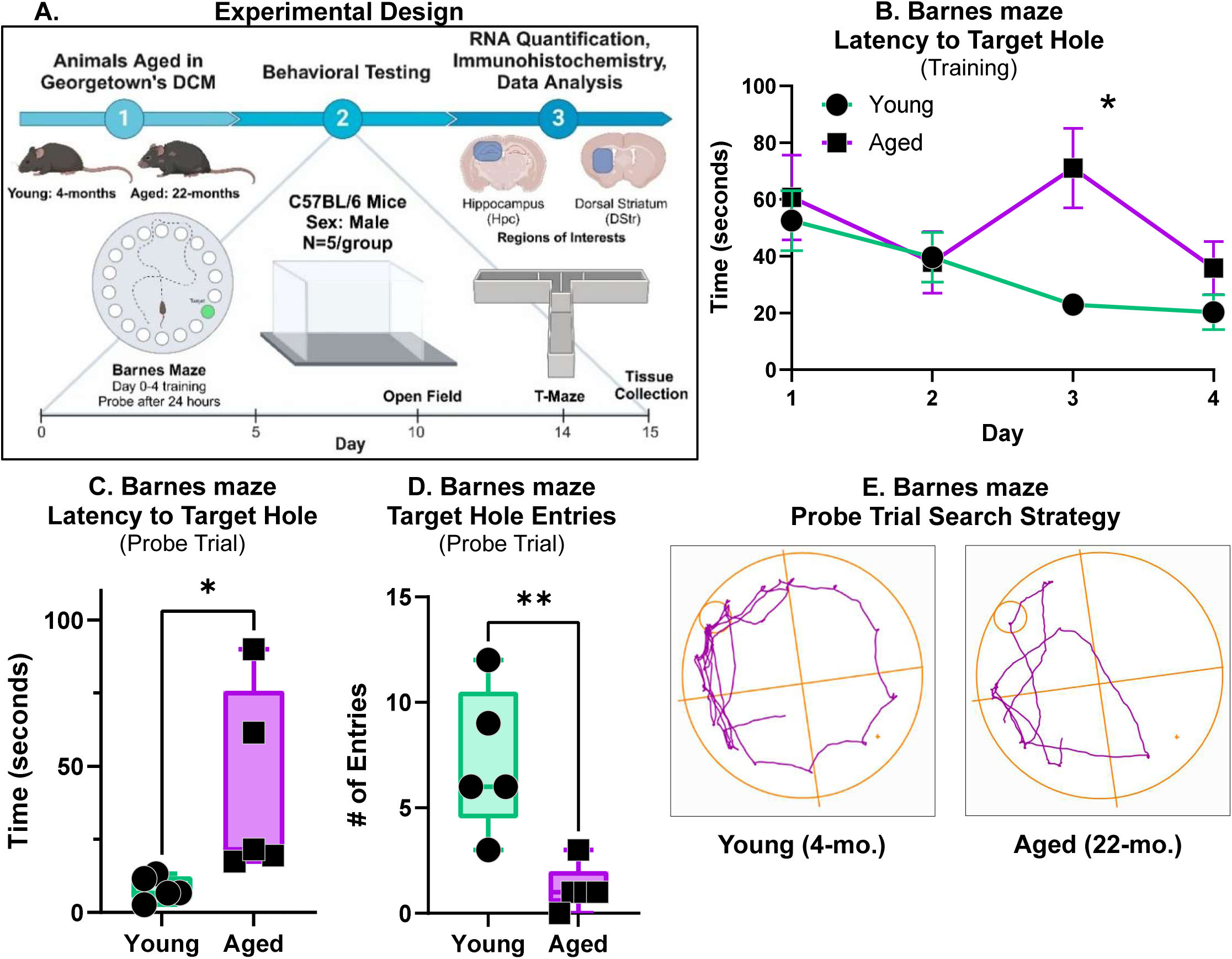
Age-related differences in spatial memory performance in the Barnes maze. **(A)** Graphic of experimental design and behavioral testing timeline. **(B)** Latency to find the target hole during the training phase, Day 1 represents the start of behavior testing. Mice underwent four days of training with three trials per day. **(C)** Latency to find target hole during the probe trial in Young and Aged mice. **(D)** Number of target hole entries during the probe trial in Young and Aged mice. **(E)** Representative images of Young (4-month-old) and Aged (22-month-old) tracking plots during probe trials. N=5 mice/group, data are presented as mean ± SEM. *p* < 0.05, **p**-**values:** *<0.05, **< 0.01. Calculated using Two-way ANOVA with Tukey or two-tailed Student t-tests.

## Results

### Aged mice have significant deficits in hippocampal-dependent learning and memory

Since cognitive deficits are implicated with aging, neurodegeneration, and changes in the extracellular matrix (1,15), a battery of behavioral tests were performed in the 14-days prior to the molecular analyses (**Fig. 1A**). To assess hippocampal-dependent visuospatial memory, we employed the Barnes maze (**Fig. 1A**). During the training phase, Aged mice learned to find the target hole at the same rate as Young mice except on Day 3 and this deficit was rescued on Day 4 of training (**Fig. 1B**). However, during the probe trial, Aged mice demonstrated a prolonged latency to locate the target hole (**Fig. 1C**) and exhibited fewer target hole entries (**Fig. 1D**) compared to Young mice. Aged mice also exhibited less efficient random search patterns to identify the missing escape hole, whereas the Young mice employed a systematic serial search strategy (**Fig 1E**).

To evaluate whether these differences were due to gross locomotor, anxiety-like, and exploratory behaviors, mice underwent the Open field behavioral test. Aged mice did not exhibit significant locomotion deficits, as indicated by the lack of change in total distance traveled and mean speed (**Sup. Fig. 1A-B**). Anxiety levels did not differ between Aged and Young mice, evidenced by no significant differences in the number of inner zone entries or the time spent in the inner zone (**Sup. Fig. 1C-D**). Additionally, mice did not show any preference for specific corners (**Sup. Fig. 1E**).

Since working memory deficits are also implicated during aging, we performed the T-maze. Aged mice did not show significant differences in the number of alternations or the latency to make a selection in both the first and second tests of each trial, suggesting that the Aged mice did not have any deficits in working memory (**Sup. Fig. 1F-G**).

### Aging affects the number of WFA^+^ and ACAN^+^ perineuronal nets and colocalization with parvalbumin cells in the hippocampus, but not in the striatum

Hippocampal-dependent memory deficits can reflect decreased plasticity within hippocampal neuronal circuits as well as deficits in the consolidation of visuospatial memory. PNNs are important for plasticity, changing in number and structure during development, in response to injury, and during aging (15,16,19). Therefore, we investigated age-related changes in PNNs, specifically PNNs surrounding PV interneurons, and asked whether there are any age-related and brain region-specific alterations in PNN and PV numbers. Following immunostaining for PV (PV^+^), *Wisteria floribunda* agglutinin (WFA^+^)-fluorescein labelling of PNNs and quantification (Hippocampus: **Fig. 2B**, Striatum: **Fig. 3B**), we found no significant differences in the total number of PV^+^ cells across the hippocampus and dorsal striatum (**Fig. 2B & 3B**, respectively). However, there was an increase in the number of double-labelled, PV^+^ WFA^+^, cells with aging in the hippocampus.

**Figure 2.**
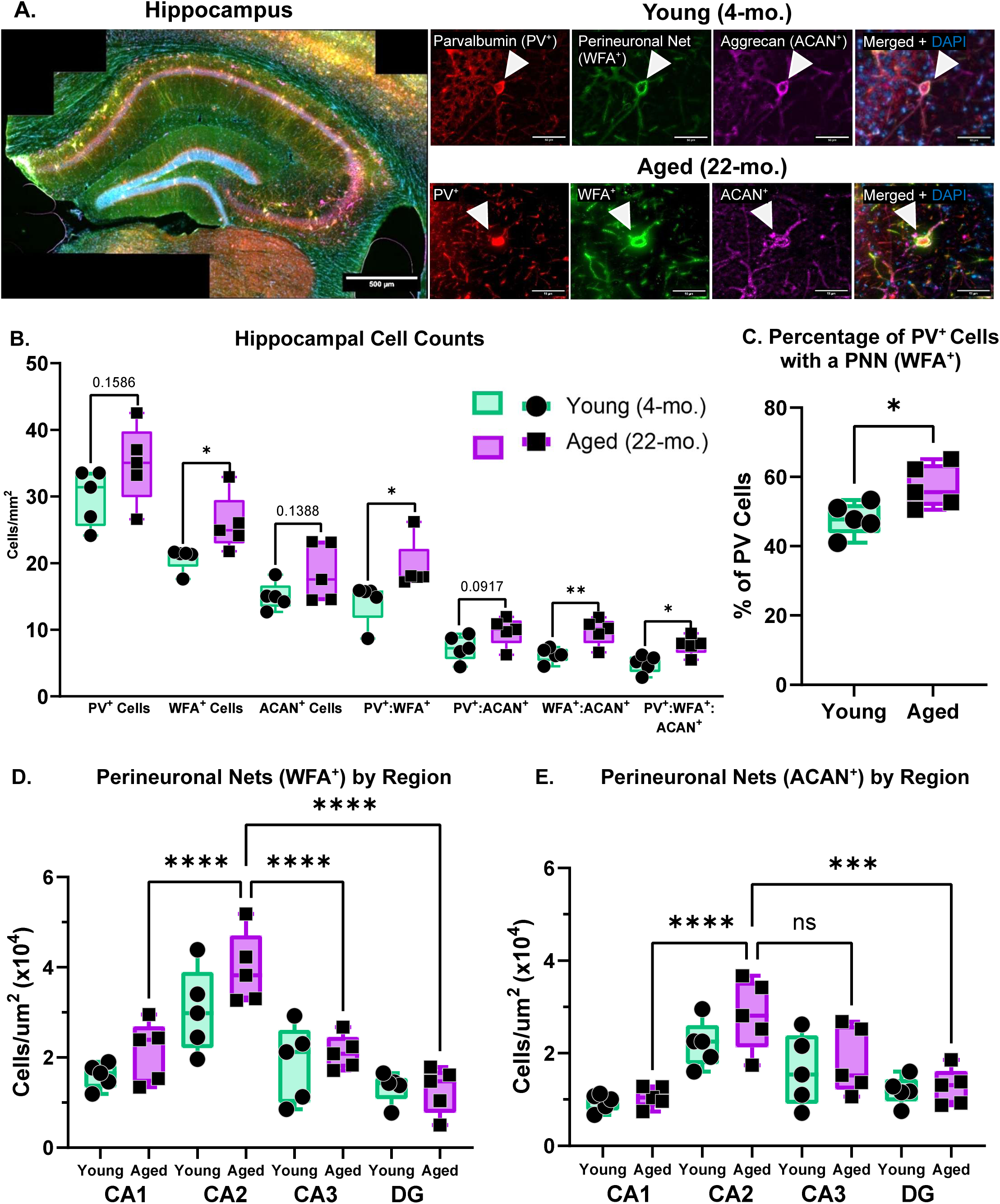
Quantification of hippocampal perineuronal nets (PNNs) and parvalbumin (PV) interneurons in Young and Aged mice. **(A)** Representative image of a stitched Aged mouse hippocampus. Representative images of Young and Aged PV^+^ interneuron, a PNN stained with *Wisteria floribunda* agglutinin (WFA^+^), a PNN stained with aggrecan (ACAN^+^), and a merged image of all three stains. **(B)** Quantification of hippocampal cell counts for PV^+^ cells, WFA^+^ PNNs, ACAN^+^ PNNs, and colocalized cell counts with PV^+^:WFA^+^, PV^+^:ACAN^+^, WFA^+^:ACAN^+^, and triple-labeled staining in Young and Aged mice. **(C)** Percentage of PV^+^ cells ensheathed by a WFA^+^ PNN. **(D-E)** Quantification of WFA^+^ and ACAN^+^ PNNs across hippocampal subregions. Only Aged regional statistical comparisons are shown on the graph. See **Supplementary Table 1** for all comparisons. N=5 mice/group, 4-8 sections/mouse, data are mean ± SEM. **p-values: \***<0.05, **< 0.01, **\*\*\***< 0.001, **\*\*\*\***< 0.0001. Calculated using Two-way ANOVA with Tukey or two-tailed Student t-tests.

**Figure 3.**
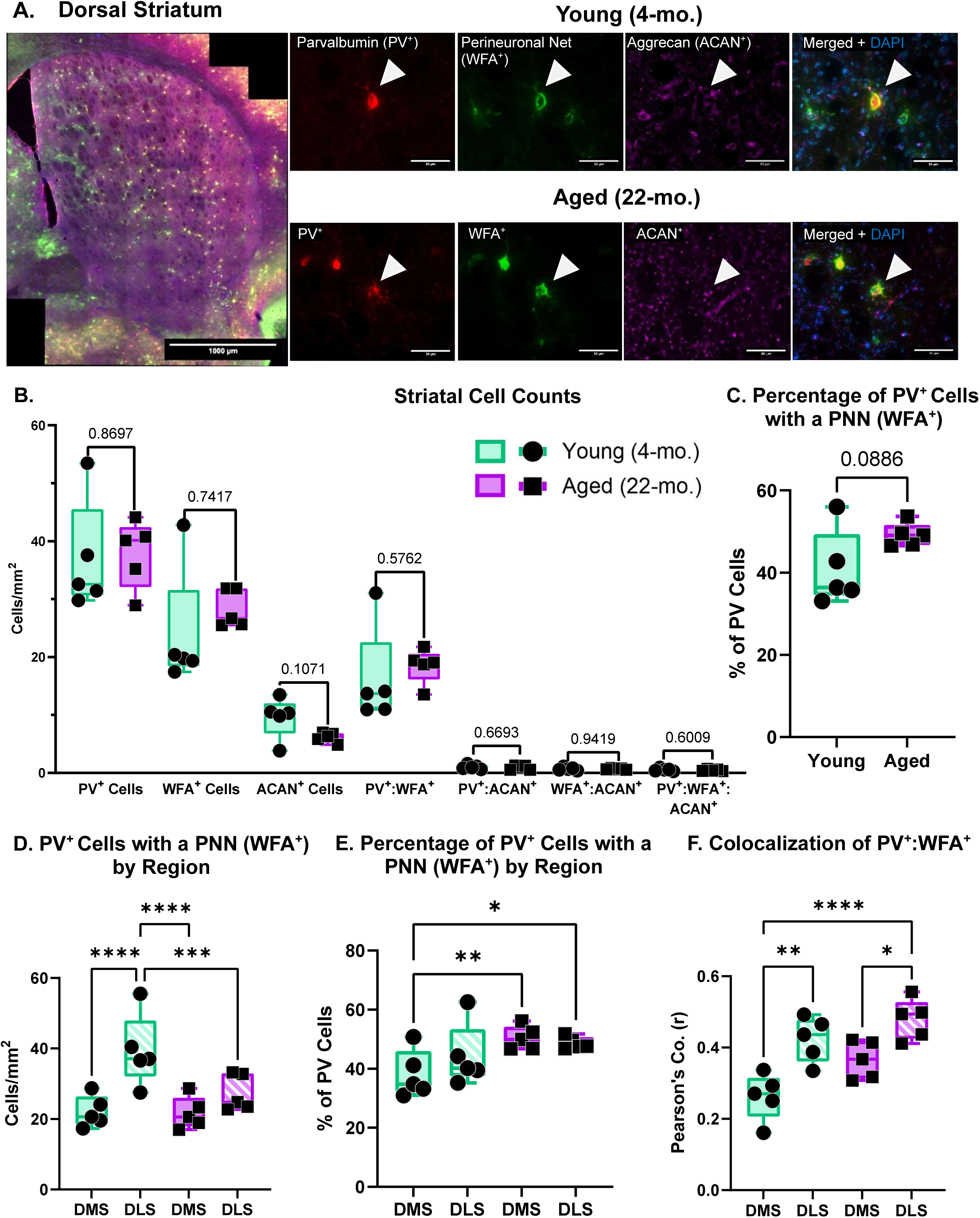
Quantification of striatal perineuronal nets and PV interneurons in Young and Aged mice. **(A)** Representative image of a stitched Aged mouse dorsal striatum, Representative images of Young and Aged PV^+^ interneuron, a PNN stained with *Wisteria floribunda* agglutinin (WFA^+^), a PNN stained with aggrecan (ACAN^+^), and a merged image of all three stains. **(B)** Quantification of cell counts for PV^+^ cells, WFA^+^ PNNs, ACAN^+^ PNNs, and colocalized cell counts with PV^+^:WFA^+^, PV^+^:ACAN^+^, WFA^+^:ACAN^+^, and triple-labeled staining in Young and Aged mice. **(C)** Percentage of PV^+^ neurons ensheathed by WFA^+^ PNNs. **(D)** Subregional quantification of co-labelled PV^+^ and WFA^+^ cells. **(E)** Subregional quantification of the percentage of PV^+^ neurons ensheathed by WFA^+^ PNNs. **(F)** Colocalization analysis of striatal subregions. Pearson’s coefficient of colocalized PV^+^:WFA^+^ neurons. N=5 mice/group, 5-8 sections/mouse, data are mean ± SEM. **p-values: \***<0.05, **< 0.01, **\*\*\***< 0.001, ****<0.0001. Calculated using Two-way ANOVA with Tukey or two-tailed Student t-tests.

Because PNNs are comprised of a variety of proteoglycans and age-related changes in plasma proteoglycans was recently reported (5), we next immunostained for aggrecan (ACAN^+^), a chondroitin sulfate proteoglycan (CSPG) expressed in the brain, and one that is important for PNN structural integrity and plasticity (35,36). In the hippocampus of Aged mice, there was a significant increase in the total number of PNNs stained with WFA and those co-stained with both WFA and aggrecan (**Fig. 2B**). There was also an increase with aging of triple-labeled cells (PV^+^:WFA^+^:ACAN^+^; **Fig. 2B**) in the hippocampus. Furthermore, the hippocampus of Aged mice had a significantly higher percentage of PV^+^ surrounded by WFA^+^ PNNs (**Fig. 2C**). When subhippocampal regions were analyzed, we found that the CA2 region had the most significant increase in the number of WFA^+^ and ACAN^+^ nets (**Fig. 2D-E**) and the highest increase in mean fluorescence intensity (**Sup. Fig. 2A, C, E** respectively) compared to the other subregions.

In contrast, there was no significant change in the total number of PNNs stained with WFA or aggrecan, nor the number of PV^+^ cells that contained a PNN in the dorsal striatum (**Fig. 3B**). Likewise, there were no significant age-related differences in total striatal PV^+^, WFA^+^, and ACAN^+^ intensities (**Sup. Fig. 2B, D, E**, respectively). There was also no difference in the percentage of PV cells that colocalized with a PNN in the total dorsal striatum (**Fig. 3C**). However, when analyzed by subregion, the Young dorsolateral striatum (DLS) had a higher number of cells co-labelled with PV^+^ and WFA^+^ compared to both the Young dorsomedial striatum (DMS) and both regions of the Aged mice (**Fig. 3D**). The Young DMS also showed a significant decrease in the percentage of PV^+^ cells with a WFA^+^ PNN when compared to both regions of the Aged striatum (**Fig. 3E**). In the colocalization analysis, the DLS showed a stronger linear relationship between WFA^+^ and PV^+^ cells compared to the DMS (**Fig. 3F**). Taken together, the Aged dorsal striatum maintained overall PNN numbers, intensity, and colocalization suggesting a maintenance of PNN homeostasis within the dorsal striatum while the hippocampus showed age-related changes.

To better understand the relationship between PV cells and PNNs, regression analyses were performed. While a significant increase in PNNs and their colocalization with PV cells was observed exclusively in the hippocampus, there was a positive linear relationship with WFA^+^ and PV^+^ in each region and age group, with a significantly strong association of WFA^+^ and PV^+^ in the Aged dorsal striatum and both age groups in the hippocampus (**Fig. 4A-B**). In addition, the Aged mice exhibited a positive association of WFA^+^ and ACAN^+^, with the hippocampus showing a significant age-related increase, compared to no observable associations in both regions of the Young mice (**Fig. 4C-D**). These results suggest that aging selectively alters PNN associations in a region-specific manner and supports that there are differential vulnerabilities of neural circuits to age-related inflammation and extracellular matrix remodeling.

**Figure 4.**
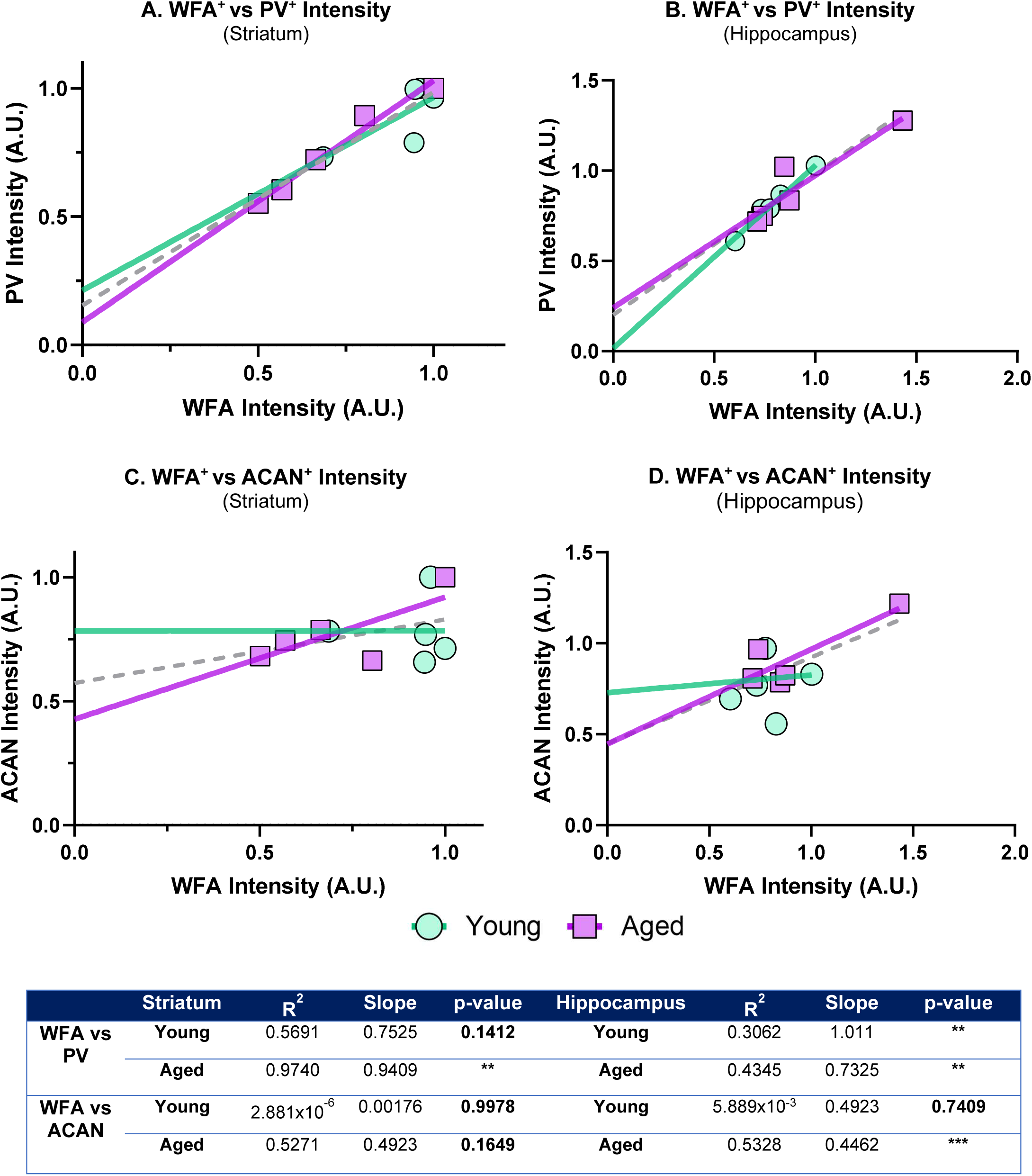
Regression analyses between WFA^+^ intensity with ACAN^+^ and PV^+^ interneurons in the hippocampus and striatum. (A–B) Linear regression analysis of WFA^+^ intensity versus PV^+^ intensity in Young and Aged mice in the striatum and hippocampus, respectively. **(C–D)** Linear regression analysis of WFA^+^ intensity versus ACAN^+^ intensity in the striatum and hippocampus, respectively. **(E)** Summary of R² values, slopes, and p-values for the coefficient differences t-test for slopes for each condition. Data was normalized before regression, and the gray dotted line represents linear regression for total samples and t-test comparison of slopes performed. **p-values:** **< 0.01, **\*\*\***< 0.001.

### Markers of gliosis, neuroinflammation, and PNNs show differential changes across brain regions with aging

We next investigated several biomarkers that impact inflammaging as well as PNN composition and homeostasis in the hippocampus and striatum of Young and Aged mice. As shown in **Figure 5**, we parsed the genes into four biomarker categories: glial, inflammatory, complement/senescence, and PNN-related markers. Overall, we noted region-specific age-related changes in gene expression for some biomarkers but not others. Both the striatum and hippocampus showed age-related increases in the astrocyte marker, glial fibrillary acidic protein (*GFAP*), and toll-like receptor 2 (*TLR2*) (**Fig. 5A**). Likewise, markers of neuroinflammation, including cytokines *IL6* and *TNFA*, chemokine *CCL2* (also known as RANTES [Regulated upon Activation, Normal T-cell Expressed and Secreted]), and complement-related genes (*C3* and *C1qa*) were significantly elevated in both brain regions in the Aged mice (**Fig. 5B-C**). P16 (*CDKN2A)*, often associated with cellular senescence was also elevated in Aged mice in both brain regions, but P21 (*CDKN1A*) did not differ between the groups (**Fig. 5C**).

**Figure 5.**
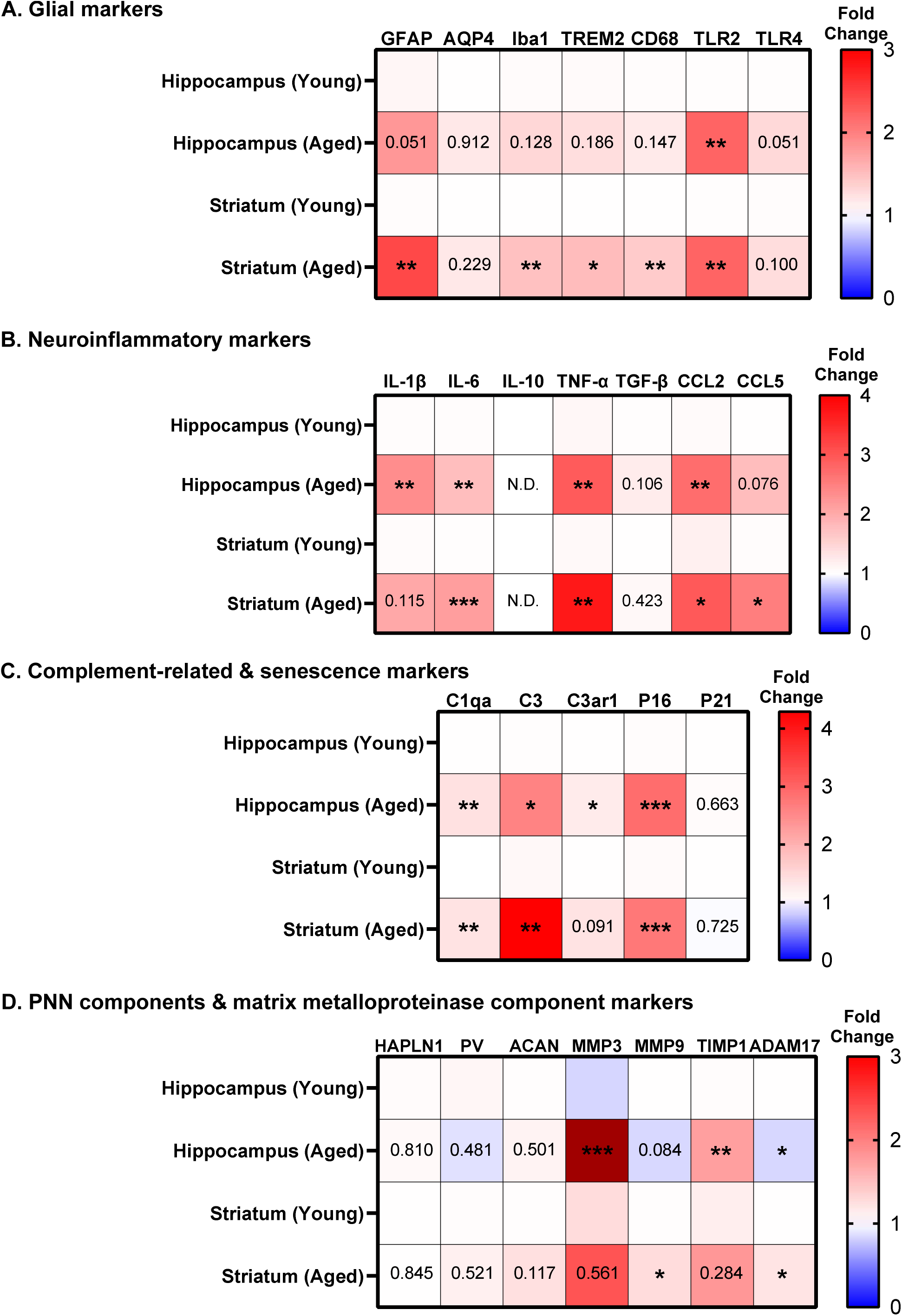
Expression of inflammatory, glial, and PNN-related markers in Aged brains. Heatmaps for expression of **(A)** Glial and pattern recognition receptor marker, **(B)** Neuroinflammatory markers, **(C)** Complement-related and senescence markers, and **(D)** PNN components and matrix metalloproteinase markers. Heat maps represent fold change relative to Young mouse counterparts; fold change is represented in the color legend to the right of the heatmap. MMP-3 dark red represents an 18.34-fold change. p-values are represented as Aged compared to Young in the corresponding marker’s Aged box. Fold changes were calculated using the ΔΔCt method, with statistical analyses performed on ΔCt values using Student t-tests. N=5 mice/group. **p-values: \***<0.05, **< 0.01, **\*\*\***< 0.001, N.D.= Not determined.

The striatum also showed region specific age-related increases in the expression of glial-associated markers: ionized calcium-binding adaptor molecule 1 (*Iba1*), triggering receptor expressed on myeloid cells-2 (*TREM2*), Cluster of Differentiation 68 (*CD68), CCL5* (Monocyte chemoattractant protein-1, MCP-1), matrix metalloproteinase 9 (*MMP9*), an inducible member of the gelatinase family, and *ADAM17*, a secreted protease that plays a role in maintenance of extracellular matrices (**Fig. 5A, B & D**). This suggests the aged striatum has elevated microgliosis and upregulation of several ECM remodeling genes. In contrast, *TLR4*, *IL1B*, C3 receptor (*C3ar1*), *MMP3* (stromelysin-1), and tissue inhibitor of metalloproteinases 1 (*TIMP1*), an endogenous inhibitor of MMPs, were all increased with aging in the hippocampus but not the striatum (**Fig. 5A-D**). The Aged hippocampus also showed a decrease in the proteases *MMP9* and *ADAM17* (**Fig. 5D**). All other markers were unchanged in both brain regions in the Aged mice compared with the Young cohort.

### Age-related glial changes reveal regional similarities and differences

We next conducted immunostaining analyses for protein markers of gliosis in Aged and Young mice. Using Iba1 to identify microglia, we did not find any difference in the total number of microglia between Aged and Young mice in the hippocampus and striatum (**Sup. Fig 3A-B**, respectively). For total mean intensity of Iba1^+^ immunostaining, the Aged striatal Iba1^+^ intensity was significantly higher than their Young counterparts (**Fig. 6B**), but no change was observed in the hippocampus (**Sup. Fig. 3C**). To further characterize microglial activation, single-cell immunostaining analysis revealed that Aged mice exhibited higher Iba1^+^ intensity per microglia compared to Young mice in both the hippocampus and the striatum, with the Aged striatum exhibiting significantly higher intensities than all other groups (**Fig. 6C**), with the Aged dorsolateral striatum showing the highest intensity per cell (**Fig. 6D**).

**Figure 6.**
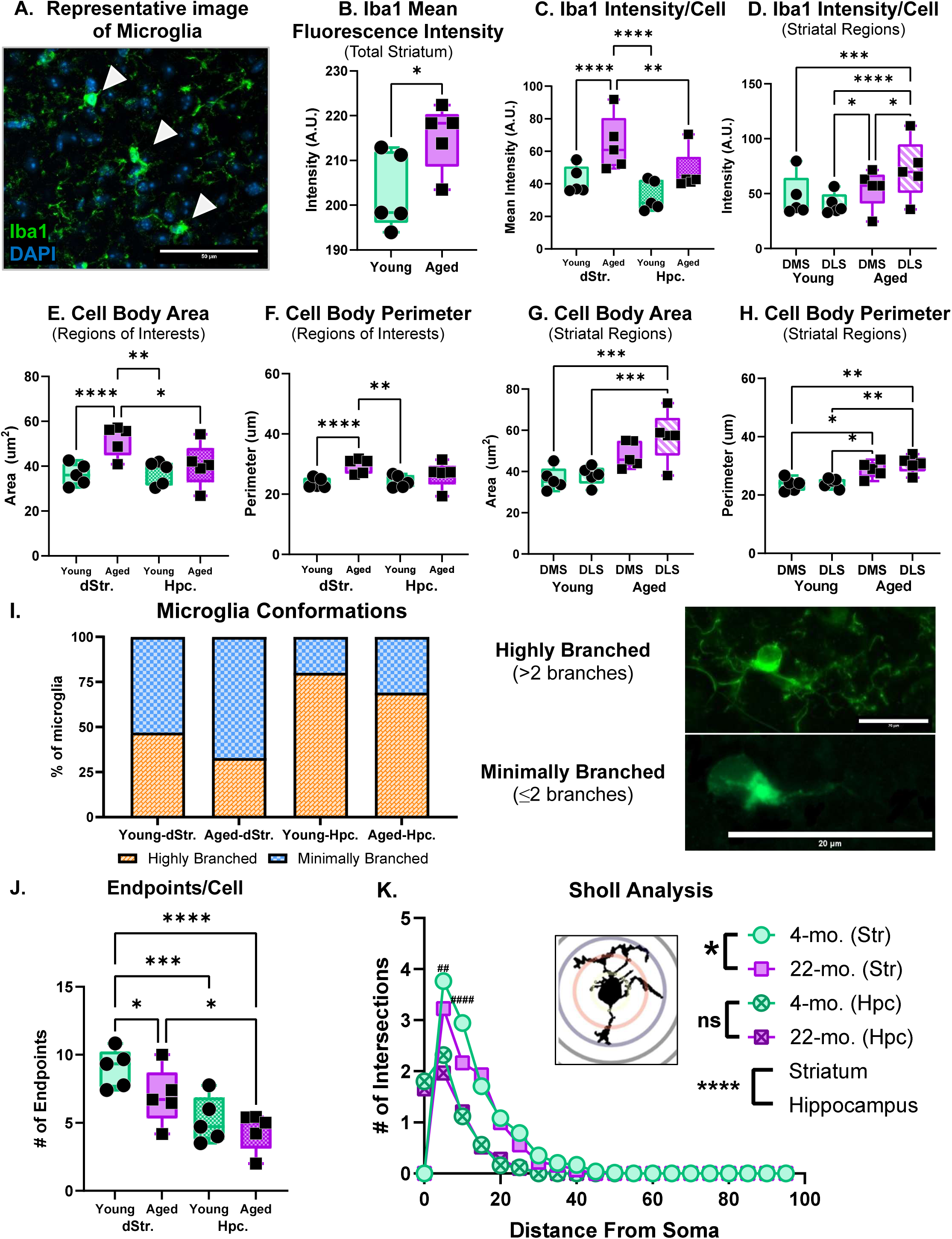
Age-related alterations in microglial activation and morphology. **(A)** Representative images of Iba1^+^ microglia in the hippocampus of an Aged mouse. **(B)** Mean Iba1^+^ fluorescence intensity for the total striatum. **(C)** Quantification of Iba1^+^ intensity per cell in the striatum and hippocampus of Young and Aged mice. **(D)** Quantification of Iba1^+^ intensity per cell by striatal subregion. **(E)** Quantification of cell body area in the striatum and hippocampus of Young and Aged mice. **(F)** Quantification of cell body perimeter in the striatum and hippocampus of Young and Aged mice. **(G-H)** Quantification of cell body area and perimeter by striatal subregions. **(I)** Categorized microglial morphological conformations across the dorsal striatum and hippocampus in Young and Aged mice with representative images of microglia classified as complex (>2 branches, ramified) or simple (≤2 branches, amoeboid-like). Data shown as percent of total microglia. Number of cells per group: dStr: Young = 64, Aged = 58; Hpc.: Young = 25, Aged = 29. **(J)** Quantification of average endpoints per cell in the striatum and hippocampus of Young and Aged mice. **(K)** Sholl analyses of the branch profile (ramification) of microglia by age and region. N=5/group with 4-14 cells/mouse. Data are mean ± SEM. **p-values: \***<0.05, **< 0.01, **\*\*\***< 0.001, ****<0.0001. Sholl Analysis striatal comparisons: **p-values:** ##<0.01, ####<0.0001. Calculated using Two-way ANOVA with Tukey or two-tailed Student t-tests. Sholl analyses were calculated using a Three-way ANOVA with Tukey.

To assess microglial morphology, which is often used as an indicator of microglial activation state (37,38), we analyzed the average cell body area and perimeter in both the striatum and hippocampus of Young and Aged mice. In the striatum, Aged microglia displayed significantly larger cell body areas (**Fig. 6E**) and perimeters (**Fig. 6F**) compared with Young mice. When analyzed regionally within the striatum, the cell body areas were significantly increased in the dorsolateral region (**Fig. 6G**) while both the dorsomedial and dorsolateral microglia in Aged mice exhibited significantly larger perimeters than both regions of the Young mice (**Fig. 6H**). To characterize microglial processes and ramifications, we characterized the branching complexity, number of endpoints, and branching patterns (i.e., Sholl profile). In terms of branching complexity, we binned individual microglia into two categories, ‘minimally branched’ and ‘highly branched.’ ‘Minimally branched’ microglia exhibited two or less branches (i.e., amoeboid-like, bipolar, or rod-like) while ‘highly branched’ microglia had greater than two branches. There are no age-related differences in characterization of microglia, however, in the striatum, the percentage of microglia characterized as ‘minimally branched’ was greater in both age groups when compared to the hippocampus (**Fig. 6I**). Endpoint analysis revealed that Aged striatal microglia had a significantly lower number of end-points per microglia compared to Young mice (**Fig. 6J**). For the Young and Aged hippocampus, there was no significant difference in the number of endpoints per microglia, and the hippocampal microglia had less endpoints per microglia than those in the striatum (**Fig. 6J**). To investigate the branching patterns we performed a Sholl analysis which revealed that striatal microglia in Young mice had significantly more intersections at 5- and 10-μm distances from the soma compared to microglia from the Aged group (**Fig 6K**). Furthermore, there was a significantly higher number of intersections at 5-25-μm distances from the soma in microglia from the striatum compared to the hippocampus (**Fig. 6K**). Overall, compared to the Young striatal microglia, the Aged microglia exhibited several morphological signs of activation including increased Iba1^+^ expression, larger cell bodies, a greater percentage of microglia characterized as ‘minimally branched’ (i.e., more amoeboid-like), fewer endpoints per microglia, and a reduced branching pattern. However, compared to the Aged hippocampus, the Aged striatal microglia also exhibited a more diverse morphology with a larger percentage of microglia exhibiting amoeboid-like morphology, while their ‘highly branched’ microglia exhibited a larger number of processes. This data suggest a possible hyper-ramified activation state in which striatal microglia exhibit increased number of branches, as well as an upregulation of immune-related genes often seen in chronic stress (39).

## Discussion

Together, our findings underscore the complex interplay between inflammation, microglial activation and perineuronal nets in aging. A more complete understanding of these interactions is crucial for developing therapeutic strategies to mitigate the effects of inflammaging and to improve cognitive health in the elderly. Recent work in the field of aging suggests that organ dysfunction occurs at different rates and that specific measures of organ biological age are important for a more comprehensive understanding of aging (reviewed in (7)). Differences in chronological and organ biological aging are likely due to the variations in genetic background and environmental factors. Here we used a rodent model which allows for control of the genetic and environmental factors as these mice were group housed and experienced the same diet and living conditions. We also focused our study on two areas of the brain that are important for cognition and motor control, the hippocampus and the striatum, which are known to be affected in the most common age-related neurodegenerative disorders, Alzheimer’s and Parkinson’s disease.

We chose behavioral and molecular measures to quantify changes between adult (Young) and old (Aged) mice. In the behavioral studies we found that Aged mice displayed significant hippocampal-dependent learning impairments in the Barnes maze, as evidenced by prolonged latencies to locate the target hole and reduced target hole entries during the probe trial, indicating differences in spatial memory retention. These hippocampal-dependent memory deficits align with the age-related increase in perineuronal nets we found in the hippocampus, suggesting that plasticity is limited in the aging hippocampus. However, Aged mice did not exhibit deficits in working memory within the T-maze paradigm and the Open field test revealed no significant age-related differences in locomotion or anxiety-like behaviors. One limitation of this study is the modest sample size which may prevented us from noting more subtle changes in these behaviors, so further investigation is warranted.

Although the observed behavioral changes were mild, the hippocampus arose as an area more vulnerable to age-related behavioral changes compared with the striatum. Since aging is also associated with immune system dysfunction, we went on to interrogate markers typical of brain inflammaging. First, we concentrated on the ECM, which is known to change in many organs during aging, is often associated with senescence, and is important for normal brain function (40). For example, in the brain, the PNN, a specialized ECM structure, helps to stabilize neurotransmitter receptors in the membrane, protects the neuron from oxidative stress and excessive calcium levels, enables cell-to-cell communication, and affects synaptic plasticity. The number and structure of the PNN influences whether plasticity can occur, such that too little or too much ECM affects neuronal function and synaptic plasticity (41). While PNNs are not evenly distributed in the brain, a majority of GABAergic parvalbumin expressing fast-spiking interneurons are surrounded by these ECM structures which consist of a hyaluronan backbone and chondroitin sulfate proteoglycans (CSPGs) including brevican, versican, neurocan and aggrecan, as well as link proteins. Here we report an increase in Aged hippocampal PNNs, which could explain the behavioral deficits observed in these mice, since an increase in the number of PNNs is linked to decreased plasticity.

Of note, alterations in PNN components, particularly the CSPGs, are associated with age-related neurodegenerative disorders such as Alzheimer’s disease and Parkinson’s disease, yet little is known about CSPG composition in normal aging. It is known that the loss of the CSPG, aggrecan, destabilizes the PNN, promoting critical period-like plasticity in the adult cortex (36) suggesting that CSPG levels are important for regulating PNN structure. Here, we show that in the aged hippocampus, there is a significant increase in total number of PNNs on PV cells. Our regression analysis revealed that while WFA^+^ intensity, a broad marker of perineuronal nets (PNNs), maintained a consistent correlation with PV⁺ intensity across age groups, Aged mice exhibited a significant increase in the association of aggrecan-containing PNNs (ACAN⁺), an important structural CSPG. This selective enrichment of ACAN⁺ nets suggest an age-dependent remodeling of PNN composition that may constrain hippocampal plasticity. Elevated aggrecan containing PNNs may restrict plasticity by impeding axonal growth required for the integration of newly formed neurons in the dentate gyrus and the neuronal connections in the hippocampal tri-synaptic circuit (36). Carstens et al. showed that removal of PNNs within CA2 restored plasticity in that region suggesting that the increase in both WFA^+^ and ACAN^+^-labelled nets shown here may reduce synaptic plasticity in aging (42). Further investigation into CSPG modifications, sulfation patterns, and the interaction between PV interneurons and hippocampal inputs will help elucidate the mechanisms underlying age-related cognitive decline. Overall, in this study we show an increase in hippocampal aggrecan positive PNNs and suggest that this finding along with our behavioral data support that there is a decrease in plasticity in the aged brain.

Other than a modest positive association with WFA^+^ and ACAN^+^ in the striatum there are no age-related changes in PNN^+^ or PV^+^ cell numbers. However, when the striatum was subdivided into the dorsolateral striatum (DLS) and the dorsomedial striatum (DMS), the DLS exhibited greater PV⁺ and WFA⁺ colocalization across both age groups. While the Young DMS showed a significant decrease in PV^+^ ensheathed by a WFA^+^ PNN. This regional distinction could reflect functional specialization. The DLS supports habit formation, sensorimotor integration, and repetitive action sequences, behaviors that benefit from stable circuitry, where PNNs may serve to reinforce input from outside the striatum. By contrast, the DMS mediates goal-directed learning and action–outcome associations, processes that demand greater synaptic flexibility. PNNs begin to form during the second postnatal week and continue to accumulate with age in the brain and the difference in the Young DMS could reflect a differential maturation of PNNs during aging in the two regions (44). This could allow for the Young mice to exhibit greater synaptic plasticity and behavioral flexibility during goal-directed learning such as during the Barnes maze, while solidifying circuits-related to habits and sensorimotor integration. Consistent with this framework, a mouse model of repetitive behaviors demonstrated an aberrant increase in PV⁺ neurons with WFA^+^-labelled PNNs within the DMS but not the DLS, and enzymatic reduction of these PNNs was sufficient to rescue the behavioral phenotype (45). Together, these findings underscore persistent regional heterogeneity that reflect functionality in striatal PNN expression. However, the positive association of WFA^+^ and ACAN^+^ and difference in the Young DMS may reflect shifts in PNN or extracellular matrix composition, warranting further investigation into the nuanced molecular changes underlying PNN remodeling in the aged brain.

PNNs can be regulated by the neuroinflammatory state of the brain since CSPGs, matrix metalloproteinases (MMPs), and enzymes that regulate MMPs are all expressed by glia and glial activation changes with aging. Our data suggests that striatal microglia, with their hyper-ramified morphology, maintain PNN homeostasis through low-level turnover facilitated by phagocytic markers (CD68) and matrix metalloproteinases (MMPs) like matrix metalloproteinase-9 (MMP-9) and A disintegrin and metalloproteinase-17 (ADAM-17). While CD68 is not exclusively expressed in microglia/macrophages, it is a lysosomal-associated membrane glycoprotein, and commonly used as a marker of phagocytic microglia/macrophages in the CNS (46). MMP-9 and ADAM-17 are both implicated in regulating microglial phagocytosis, likely through extracellular matrix remodeling which supports debris clearance and cell migration (47–49). In contrast, the hippocampus exhibited increased PNN counts, WFA^+^ intensity, and aggrecan colocalization in Aged mice, suggesting a less plastic environment contributing to hippocampal-dependent spatial memory deficits. MMP-3 is also critical for extracellular matrix remodeling and was upregulated in the hippocampus, however, TIMP-1, an endogenous MMP inhibitor was also increased. Of note, MMP-9 and ADAM-17 were also decreased in the hippocampus which we hypothesize is reducing overall matrix turnover and limiting hippocampal plasticity.

How the PNNs are regulated is an important question and here we showed that Aged mice had a significant increase in markers of neuroinflammation, complement activation, and cellular senescence, with clear regional specificity suggesting a role for inflammaging in the normal aging process. Notably, IL-6 and TNF-α, two key pro-inflammatory cytokines associated with inflammaging, were elevated in Aged brains. These molecules are known to disrupt the blood-brain barrier (BBB), activate microglia into neurotoxic states, and impair both synaptic plasticity and neurogenesis (4). Chronic IL-6 and TNF-α have been linked to cognitive decline and neurodegenerative diseases with their levels correlating to disease severity (50–53). In the hippocampus, IL-6 interferes with long-term potentiation (54), while TNF-α impacts synaptic strength and dendritic spine density, contributing to reduced hippocampal volume and memory deficits (55). Additionally, elevated IL-1β in the hippocampus is associated with amyloid and tau pathology, as it promotes neuroinflammation, disrupts synaptic function, and inhibits neurogenesis in AD patients (56–58).

Additionally, we noted an upregulation in TLR-2 in Aged mice, which following activation can create a feedback loop amplifying inflammation in the brain (reviewed in (59)). TLR-2 is expressed by neurons, microglia, astrocytes, and peripheral immune cells, and triggers the NF-κB signaling pathway, promoting the release of inflammatory molecules like IL-6, TNF-α, and complement proteins (60), which we also show are increased in the aged brain. NF-κB signaling also regulates chemokine expression (i.e., CCL-2 and CCL-5) further exacerbating neuroinflammation by recruiting immune cells to the central nervous system, activating microglia, and disrupting the BBB (61). These chemokines are also part of the senescence-associated secretory phenotype, linking them to cellular senescence pathways (62). The elevated P16 levels in the Aged mice suggest that further research is needed to determine whether microglia undergo senescence and how these inflammatory molecules tie into senescence, but this is a plausible scenario since aging is associated with the increased numbers of senescence cells (62) and is supported by our findings.

An interesting aspect of this study is the increased expression of innate immune markers, particularly complement proteins C3 and C1qa in Aged mice. These proteins, essential for synaptic pruning, debris clearance, and microglial activation, are elevated in aged mice (63). Chronic elevations of C1qa and C3 are linked to inflammaging, though further research is needed to determine whether their role in aging is protective or detrimental. Overall, our findings underscore the intricate role of innate immune processes in aging and neuroinflammation.

This study identifies several possible glial pathways that can affect PNN homeostasis. Morphological analysis revealed that Aged microglia in the dorsal striatum shows signs of activation, including increased expression of Iba1, TREM2, CD68, greater Iba1^+^ intensity, larger cell bodies, and greater perimeter measurements compared to their Young counterparts. Microglial morphology is linked to their activation state, with ramified, highly branched processes characteristic of surveillant, homeostatic microglia, and retracted, amoeboid shapes indicating a reactive, pro-inflammatory phenotype (37,39). Transitions in morphology often reflect functional shifts, such as increased phagocytic activity or cytokine release in response to injury or disease (38). Microglia are dynamic in their ability to switch between phenotypes in response to the environment, aging, or disease-causing pathology (37–39). There are also subcategories of microglia within different morphological phenotypes (64). Here we report microglial morphology differences between Young and Aged mice as well as regional differences. In our study, Aged striatal microglia had larger cell bodies and a greater proportion of ‘minimally branched’ microglia. However, the highly branched striatal microglia had more ramifications (in endpoint and Sholl analysis) compared to Aged hippocampal microglia, we suggest this hyper-ramified morphology helps maintain the extracellular matrix homeostasis we noted in the striatum. Hyper-ramified microglia are thought to be a functional intermediate or primed state of microglia activation (65) and possibly serve as an adaptive, synapse-modulating phenotype under stress conditions that could lead to neuronal circuit resilience (66). These microglia may be important drivers in the maintenance of PNN homeostasis within the striatum during inflammaging-related conditions. The age-related decrease in endpoints and number of intersections in the Aged striatum does suggest an age-associated change in microglia morphology that may reflect a possible vulnerability during aging. We also found differential gene expressions of microgliosis and phagocytosis markers between the striatum and hippocampus. Specifically, the gene expression for markers such as Iba1, TREM2, and CD68 were significantly elevated in the striatum but not in the hippocampus. This along with increased Iba1 protein expression in the Aged striatum and gene expression of the proteases MMP-9 and ADAM-17, which are involved in the activation of inflammatory molecules and extracellular matrix remodeling, could indicate a higher level of microglial activity in the striatum. These findings suggest a heightened ability of striatal microglia to dynamically regulate PNNs in normal aging compared to hippocampal microglia. As mentioned previously, the maintenance of PNN plasticity requires a fine balance of PNN, such that too little or too much deposition can alter neuronal function.

Taken together, these findings highlight the complex interplay between neuroinflammation, immune signaling, and PNNs during normal aging, underscoring the need for further investigation into their roles in age-related cognitive decline and neurodegeneration.

## Limitations and future prospects

This study has several limitations. Only male mice were utilized in this study; future research should include sex as a biological variable to provide a more comprehensive understanding. The sample size in this study was modest yet sufficient to detect significant differences related to aging, though more subtle effects may have been masked due to the study’s statistical power. While we approximated the human-equivalent age in mice, factors such as environmental influences, dietary variations, and genetic diversity that impact human aging were not fully captured in wild-type mice housed in controlled conditions designed to minimize confounding variables. Future studies should address gene expression analyses at the single-cell level to better determine the role of each cell type in aging.

## Conclusion

In conclusion, our study shows that aged mice exhibit significant impairments in hippocampal-dependent learning, aligning with increased PNN deposition. Changes in pro-inflammatory markers, MMPs, and markers of gliosis suggest that microglial activation influences PNN homeostasis in an age and brain region dependent manner. The striatum maintained PNN integrity, whereas the hippocampus showed increased PNN counts, which we suggest could lead to impaired spatial memory through decreased plasticity. These findings suggest that aging is characterized by region-specific changes in neuroinflammation, glial activation, and PNN homeostasis, which together may underlie the cognitive decline often observed in older individuals. This work also supports that further research is needed to explore therapeutic strategies targeting inflammaging.

## List of Abbreviations

Aβ: Amyloid-beta
ACAN: Aggrecan
AD: Alzheimer’s disease
ADAM: A disintegrin and metalloproteinase
BBB: Blood-brain barrier
C1qa: Complement component 1, q subcomponent, A chain
C3: Complement component 3
C3ar1: Complement component 3a receptor 1
CA1: Cornu Ammonis field 1
CA2: Cornu Ammonis field 2
CA3: Cornu Ammonis field 3
CCL2/MCP-1: Chemokine (C-C motif) ligand 2 or Monocyte chemoattractant protein-1
CCL5/RANTES: Chemokine (C-C motif) ligand 5 or Regulated on activation, normal T-cell expressed and secreted
CD68: Cluster of differentiation 68
CSPG: Chondroitin sulfate proteoglycan
DG: Dentate gyrus
DLS: Dorsolateral striatum
DMS: Dorsomedial striatum
Iba1: Ionizing calcium-binding adaptor molecule-1
IL-6: Interleukin
MMP: Matrix metalloproteinases
NF-κB: Nuclear factor kappa-light-chain-enhancer of activated B-cells
PD: Parkinson’s disease
PNN: Perineuronal nets
PV: Parvalbumin
ROS: Reactive oxygen species
TIMP1: Tissue inhibitor of metalloproteinases 1
TLR: Toll-like receptors
TNF-α: Tumor necrosis factor alpha
TREM2: Triggering receptor expressed on myeloid cells 2
WFA: Wisteria floribunda agglutinin

## Acknowledgements

We acknowledge and thank Gianna Mamone, Javada Faison, and Fenn Suter for their assistance with sectioning tissues and blinded cell counting analyses. We would like to acknowledge Scott Litwiler for his staining and analysis of microglia cell counts; the Georgetown University Department of Neuroscience behavioral and microscopy facilities were used during the study. We would like to express our gratitude to the Georgetown University Division of Comparative Medicine veterinarians and veterinary technicians for the care of the mice.

## Author contributions

ZAC and KM designing and conceptualization, ZAC performed the majority of the experiments with assistance from SC and SL for the microglia experiments. GM, JF, FS assisted with tissue processing and blinded cell counts for the PNN experiments. ZAC analyzed, graphed, and wrote the majority of the first draft of the manuscript with the assistance of SC in writing of methods and results for microglia cell morphology experiments. ZC and KMZ wrote the revisions. All authors read, reviewed, suggested improvements, and approved the manuscript.

## Data availability

Data supplied in **Supplementary Table 1 and 2** for all experiments listed in the figures. All other data will be provided upon reasonable request from the corresponding author.

## Ethics approval

All procedures and experiments were approved by Georgetown University’s Institutional Animal Care and Use Committee and followed the United States’ National Institute of Health ethical guidelines. Animals were housed in Georgetown University Division of Comparative Medicine facilities with ad libitum water and food in a 12H light/dark cycle. Animals were anesthetized and checked for reaction to aversive stimuli via tail pinch prior to perfusion.

## Author detail

Zachary A. Colon is a fourth-year doctoral candidate in the Interdisciplinary Program in Neuroscience (IPN) in Georgetown University’s Biomedical Graduate Education, with a Master’s in Integrative Neuroscience (Georgetown University) and a Bachelor of Science in Microbiology (University of Washington). Shannon C. Chan is a graduate of Georgetown University College of Arts & Sciences’ Department of Biology with a Bachelor of Science in Neurobiology. Kathleen A. Maguire-Zeiss is a professor and Chair of the Department of Neuroscience at Georgetown University School of Medicine, and the Director of the Interdisciplinary PhD Program in Neuroscience (IPN).

