## Supplementary Figures for "Age-related inflammatory changes and perineuronal net dynamics: implications for neurodegenerative disorders"

### Supplemental Figure 1:

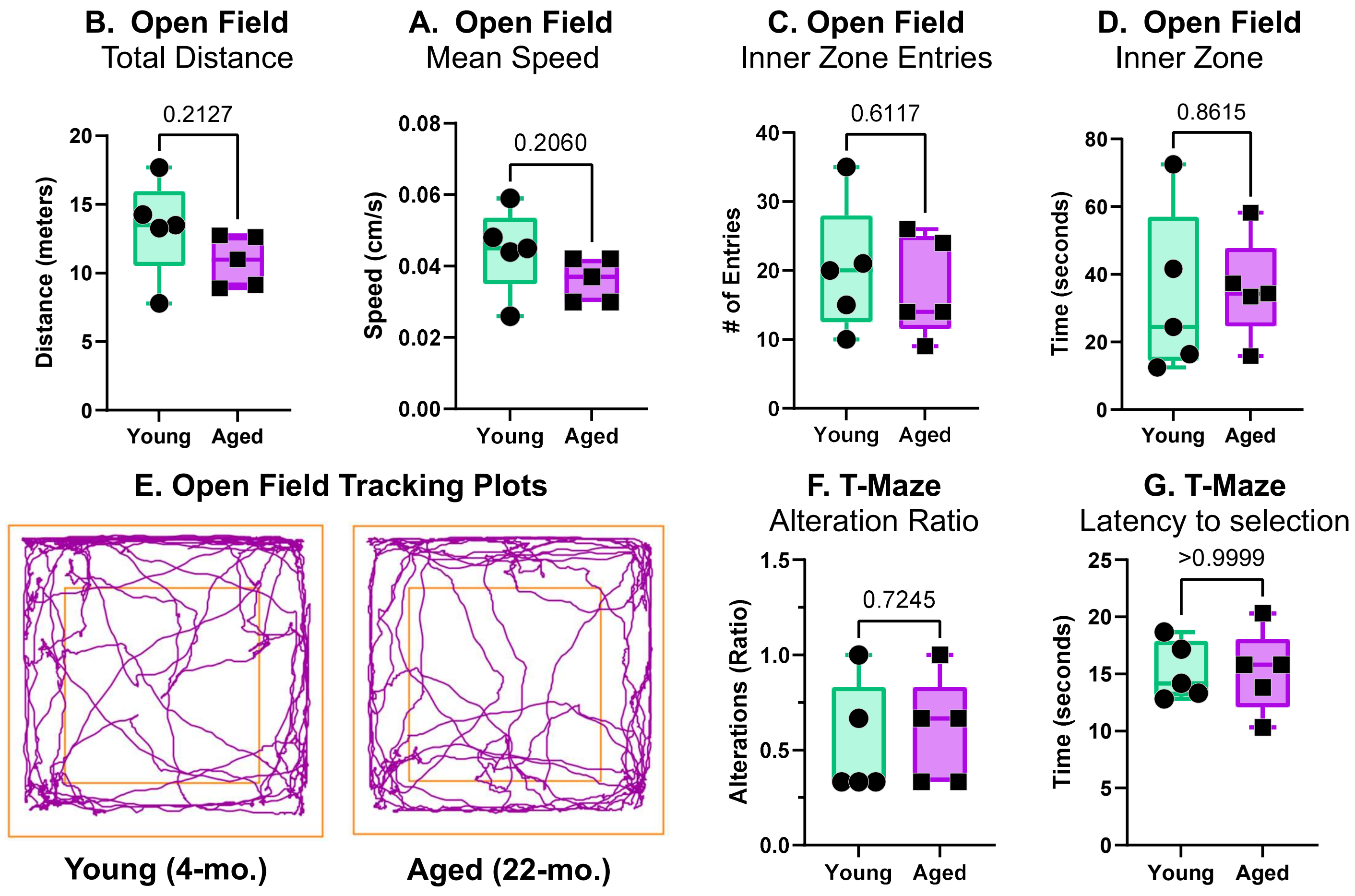


**<u>Supplemental Figure 1 Legend:</u> Open field and T-Maze behavioral performance in Young and Aged mice. (A)** Total distance travelled during Open field trial. **(B)** Mean speed of mice during Open field trial. **(C)** Number of entries into the inner zone of Open field apparatus. **(D)** Total time spent within the inner zone during Open field trial. **(E)** Representative tracking plots of mice during Open field trial. **(F)** Ratio of alterations made during second run of each T-Maze trial. **(G)** Latency to make selection per T-Maze trial. N=5 mice/group, data are mean ± SEM. p-value set to <0.05. Calculated using two-tailed Student t-tests.

### Supplemental Figure 2:

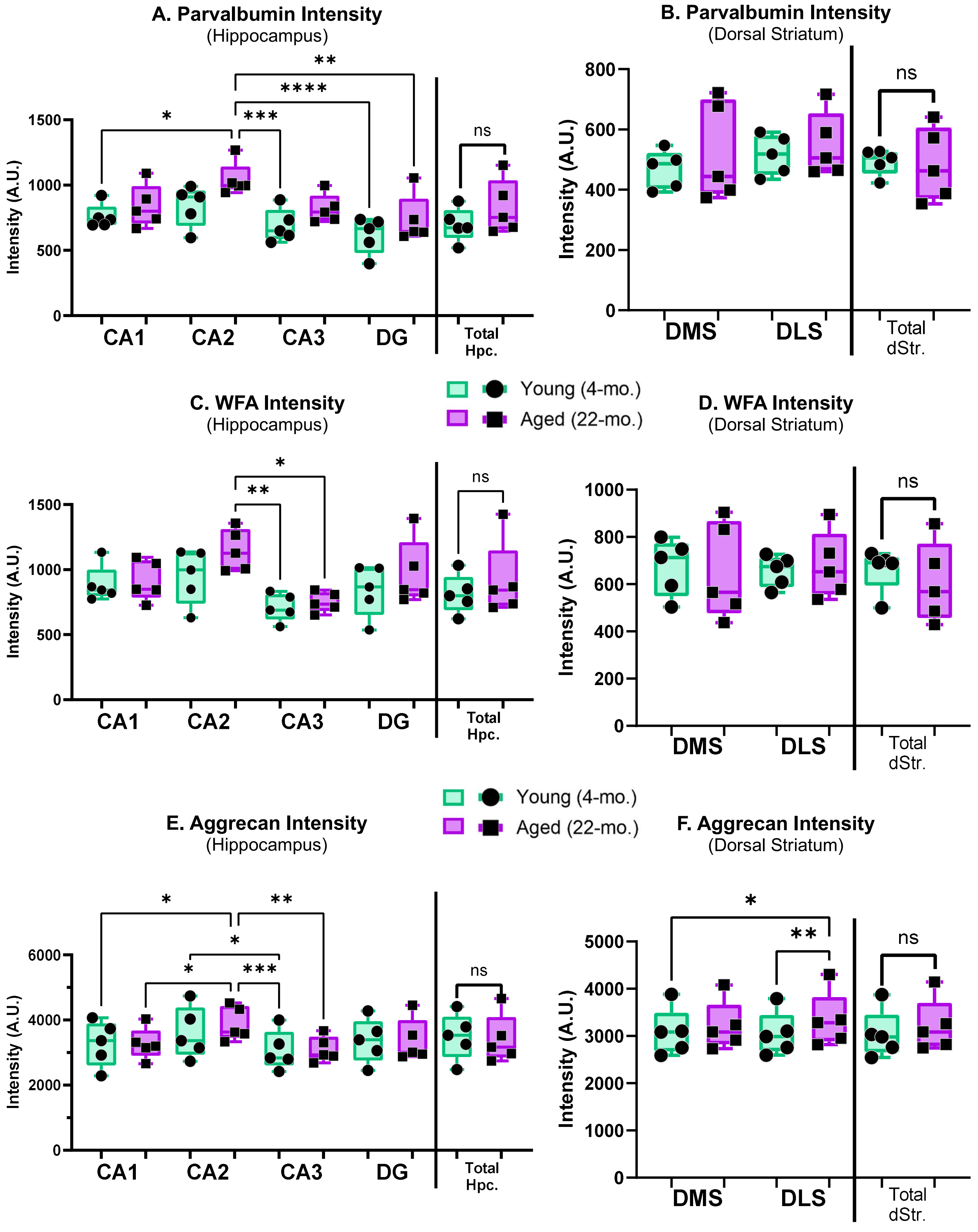

**Supplemental Figure 2 Legend:** **Mean intensities by hippocampal and striatal regions. (A)** Hippocampal parvalbumin ($PV^+$) intensity by region and total hippocampus. **(B)** Striatal $PV^+$ intensity by region and total striatum. **(C)** Hippocampal *Wisteria floribunda* agglutinin ($WFA^+$) intensity by region and total hippocampus. **(D)** Striatal $WFA^+$ intensity by region and total striatum. **(E)** Hippocampal aggrecan ($ACAN^+$) intensity by region and total hippocampus. **(F)** Striatal $ACAN^+$ intensity by region and total striatum. N=5 mice/group, 4-8 sections/mouse, data are mean ± SEM. **p-values:** *<0.05, **< 0.01, ***< 0.001, ****< 0.0001. Calculated using Two-way ANOVA w/ Tukey or two-tailed Student t-tests.

### Supplemental Figure 3:

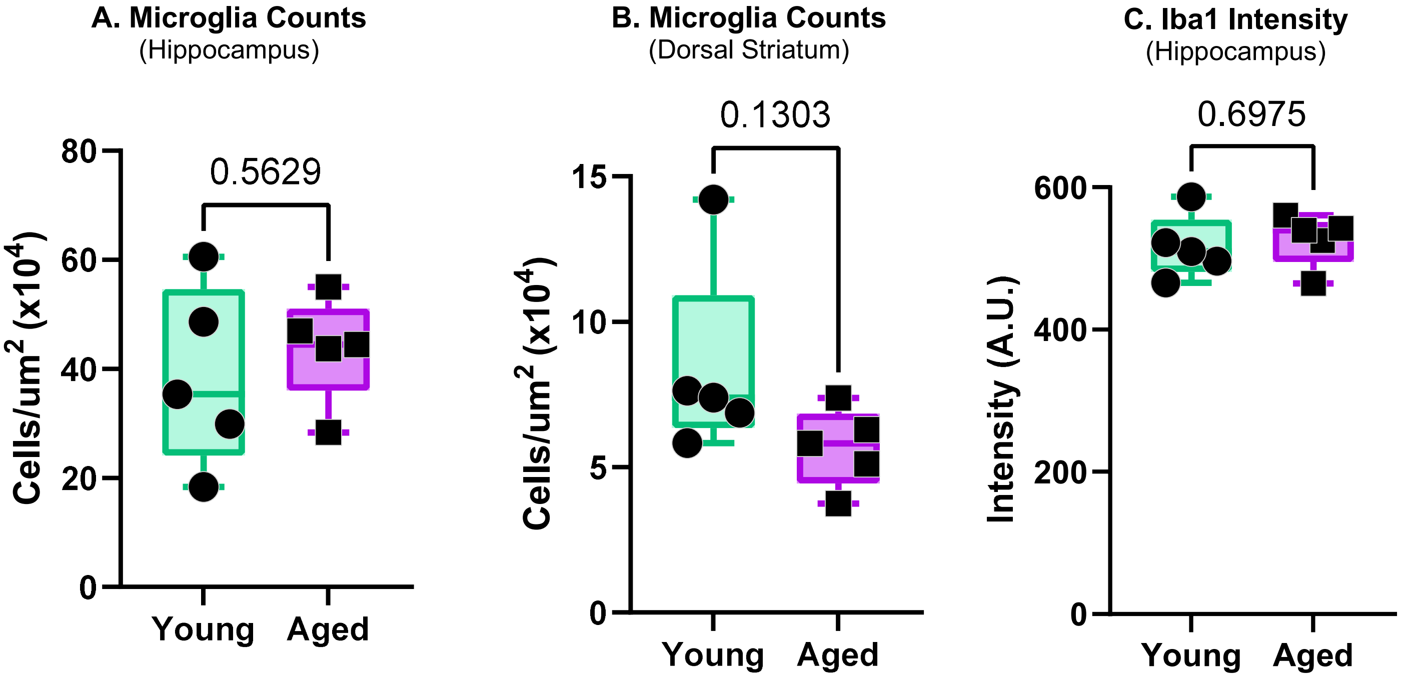


**Supplemental Figure 3 Legend:** **Microglial cell counts and hippocampal $Iba1^+$ intensity. (A)** Total microglial counts in the hippocampus. **(B)** Total microglial counts in the dorsal striatum. **(C)** Total Mean $Iba1^+$ intensity in the hippocampus. N=5 mice/group, 4-8 sections/mouse, data are mean ± SEM. p-value set to <0.05. Calculated using two-tailed Student t-tests.
