## Supplementary Tables for "Age-related inflammatory changes and perineuronal net dynamics: implications for neurodegenerative disorders"

### Supplementary Table 1

| Figure 1 | Subfigure | Experiment | Strain | Young | Aged | Statistical test | Day/Trial | Young Mean | SEM | Aged Mean | SEM | Units | p-value | Significance |
|---|---|---|---|---|---|---|---|---|---|---|---|---|---|---|
| 1 | B | Barnes maze-Latency to find target hole (Training) | C57/BL6 | N=5 | N=5 | Two-way ANOVA w/ Tukey | 0 | 52.56833 | 10.65164 | 60.74167 | 14.97003 | seconds | 0.6694 | ns |
| | | | | | | | 1 | 39.665 | 8.752984 | 37.915 | 10.89164 | seconds | 0.9036 | ns |
| | | | | | | | 2 | 22.99 | 2.62284 | 71.065 | 14.01323 | seconds | 0.0252 | * |
| | | | | | | | 3 | 20.255 | 6.099116 | 35.91 | 9.389098 | seconds | 0.2055 | ns |
| | | | | | | | Group | 33.87 | | 51.41 | | seconds | 0.1021 | ns |
| | C | Barnes maze-Latency to find target hole (Probe trail) | C57/BL6 | N=5 | N=5 | Student t-test | Probe | 8.16 | 1.892 | 42.02 | 14.5 | seconds | 0.0493 | * |
| | D | Barnes maze-Target hole entries (Probe trial) | C57/BL6 | N=5 | N=5 | Student t-test | Probe | 7.2 | 1.53 | 1.2 | 0.4899 | # of entries | 0.0057 | ** |
| **Figure 2** | **Subfigure** | **Experiment** | **Strain** | **Young** | **Aged** | **Statistical test** | **Comparison** | **Young Mean** | **SEM** | **Aged Mean** | **SEM** | **Units** | **p-value** | **Significance** |
| 2 | B | Hippocampal cell counts | C57/BL6 | N=5, 2-7 sections/ mouse | N=5, 3-4 sections/ mouse | Student t-test | PV$^+$ Cells | 29.91 | 1.874 | 34.88 | 2.595 | # of cells/mm$^2$ | 0.1586 | ns |
| | | | | | | | WFA$^+$ Cells | 20.69 | 0.7683 | 25.96 | 1.876 | # of cells/mm$^2$ | 0.0318 | * |
| | | | | | | | ACAN$^+$ Cells | 15.04 | 0.9128 | 18.58 | 1.95 | # of cells/mm$^2$ | 0.1388 | ns |
| | | | | | | | PV$^+$:WFA$^+$ Cells | 14.19 | 1.386 | 19.47 | 1.69 | # of cells/mm$^2$ | 0.0421 | * |
| | | | | | | | PV$^+$:ACAN$^+$ Cells | 7.308 | 0.8589 | 9.777 | 0.9607 | # of cells/mm$^2$ | 0.0917 | ns |
| | | | | | | | WFA$^+$:ACAN$^+$ Cells | 6.248 | 0.4854 | 9.77 | 0.8901 | # of cells/mm$^2$ | 0.0084 | ** |
| | | | | | | | PV$^+$:WFA$^+$:ACAN$^+$ Cells | 4.907 | 0.5917 | 7.696 | 0.6379 | # of cells/mm$^2$ | 0.0125 | * |
| | C | Percentage of PV$^+$ Cells with a WFA$^+$ PNN (Hippocampus) | C57/BL6 | N=5, 2-7 sections/ mouse | N=5, 3-4 sections/ mouse | Student t-test | Young vs Aged | 47.95 | 2.102 | 57.27 | 2.783 | % of PV cells | 0.0282 | * |
| | | | | | | | **[A] vs [B]** | **[A] Mean** | **[A] SEM** | **[B] Mean** | **[B] SEM** | **Units** | **p-value** | **Significance** |
| | D | WFA$^+$ PNNs by hippocampal subregion | C57/BL6 | N=5, 2-7 sections/ mouse | N=5, 3-4 sections/ mouse | Two-way ANOVA w/ Tukey | CA1 (Young) vs. CA1 (Aged) | 0.0001594 | 0.00001243 | 0.0002128 | 0.00003012 | # of cells/μm$^2$ | 0.7598 | ns |
| | | | | | | | CA1 (Young) vs. CA2 (Young) | 0.0001594 | 0.00001243 | 0.0003036 | 0.00004165 | # of cells/μm$^2$ | 0.001 | *** |
| | | | | | | | CA1 (Young) vs. CA2 (Aged) | 0.0001594 | 0.00001243 | 0.000396 | 0.0000353 | # of cells/μm$^2$ | <0.0001 | **** |
| | | | | | | | CA1 (Young) vs. CA3 (Young) | 0.0001594 | 0.00001243 | 0.0001864 | 0.00003831 | # of cells/μm$^2$ | 0.9926 | ns |
| | | | | | | | CA1 (Young) vs. CA3 (Aged) | 0.0001594 | 0.00001243 | 0.0002106 | 0.00001688 | # of cells/μm$^2$ | 0.7967 | ns |
| | | | | | | | CA1 (Young) vs. DG (Young) | 0.0001594 | 0.00001243 | 0.0001336 | 0.00001486 | # of cells/μm$^2$ | 0.9944 | ns |
| | | | | | | | CA1 (Young) vs. DG (Aged) | 0.0001594 | 0.00001243 | 0.0001283 | 0.00002307 | # of cells/μm$^2$ | 0.9833 | ns |
| | | | | | | | CA1 (Aged) vs. CA2 (Young) | 0.0002128 | 0.00003012 | 0.0003036 | 0.00004165 | # of cells/μm$^2$ | 0.1364 | ns |
| | | | | | | | CA1 (Aged) vs. CA2 (Aged) | 0.0002128 | 0.00003012 | 0.000396 | 0.0000353 | # of cells/μm$^2$ | <0.0001 | **** |
| | | | | | | | CA1 (Aged) vs. CA3 (Young) | 0.0002128 | 0.00003012 | 0.0001864 | 0.00003831 | # of cells/μm$^2$ | 0.9938 | ns |
| | | | | | | | CA1 (Aged) vs. CA3 (Aged) | 0.0002128 | 0.00003012 | 0.0002106 | 0.00001688 | # of cells/μm$^2$ | >0.9999 | ns |
| | | | | | | | CA1 (Aged) vs. DG (Young) | 0.0002128 | 0.00003012 | 0.0001336 | 0.00001486 | # of cells/μm$^2$ | 0.2785 | ns |
| | | | | | | | CA1 (Aged) vs. DG (Aged) | 0.0002128 | 0.00003012 | 0.0001283 | 0.00002307 | # of cells/μm$^2$ | 0.2086 | ns |
| | | | | | | | CA2 (Young) vs. CA2 (Aged) | 0.0003036 | 0.00004165 | 0.000396 | 0.0000353 | # of cells/μm$^2$ | 0.1222 | ns |
| | | | | | | | CA2 (Young) vs. CA3 (Young) | 0.0003036 | 0.00004165 | 0.0001864 | 0.00003831 | # of cells/μm$^2$ | 0.0156 | * |
| | | | | | | | CA2 (Young) vs. CA3 (Aged) | 0.0003036 | 0.00004165 | 0.0002106 | 0.00001688 | # of cells/μm$^2$ | 0.1172 | ns |
| | | | | | | | CA2 (Young) vs. DG (Young) | 0.0003036 | 0.00004165 | 0.0001336 | 0.00001486 | # of cells/μm$^2$ | <0.0001 | **** |
| | | | | | | | CA2 (Young) vs. DG (Aged) | 0.0003036 | 0.00004165 | 0.0001283 | 0.00002307 | # of cells/μm$^2$ | <0.0001 | **** |
| | | | | | | | CA2 (Aged) vs. CA3 (Young) | 0.000396 | 0.0000353 | 0.0001864 | 0.00003831 | # of cells/μm$^2$ | <0.0001 | **** |
| | | | | | | | CA2 (Aged) vs. CA3 (Aged) | 0.000396 | 0.0000353 | 0.0002106 | 0.00001688 | # of cells/μm$^2$ | <0.0001 | **** |
| | | | | | | | CA2 (Aged) vs. DG (Young) | 0.000396 | 0.0000353 | 0.0001336 | 0.00001486 | # of cells/μm$^2$ | <0.0001 | **** |
| | | | | | | | CA2 (Aged) vs. DG (Aged) | 0.000396 | 0.0000353 | 0.0001283 | 0.00002307 | # of cells/μm$^2$ | <0.0001 | **** |
| | | | | | | | CA3 (Young) vs. CA3 (Aged) | 0.0001864 | 0.00003831 | 0.0002106 | 0.00001688 | # of cells/μm$^2$ | 0.9964 | ns |
| | | | | | | | CA3 (Young) vs. DG (Young) | 0.0001864 | 0.00003831 | 0.0001336 | 0.00001486 | # of cells/μm$^2$ | 0.7665 | ns |
| | | | | | | | CA3 (Young) vs. DG (Aged) | 0.0001864 | 0.00003831 | 0.0001283 | 0.00002307 | # of cells/μm$^2$ | 0.6725 | ns |
| | | | | | | | CA3 (Aged) vs. DG (Young) | 0.0002106 | 0.00001688 | 0.0001336 | 0.00001486 | # of cells/μm$^2$ | 0.3133 | ns |
| | | | | | | | CA3 (Aged) vs. DG (Aged) | 0.0002106 | 0.00001688 | 0.0001283 | 0.00002307 | # of cells/μm$^2$ | 0.2376 | ns |
| | | | | | | | DG (Young) vs. DG (Aged) | 0.0001336 | 0.00001486 | 0.0001283 | 0.00002307 | # of cells/μm$^2$ | >0.9999 | ns |
| | E | ACAN$^+$ PNNs by hippocampal subregion | C57/BL6 | N=5, 2-7 sections/ mouse | N=5, 3-4 sections/ mouse | Two-way ANOVA w/ Tukey | CA1 (Young) vs. CA1 (Aged) | 0.0000944 | 0.000008325 | 0.0001058 | 0.00001013 | # of cells/μm$^2$ | >0.9999 | ns |
| | | | | | | | CA1 (Young) vs. CA2 (Young) | 0.0000944 | 0.000008325 | 0.0002196 | 0.00002244 | # of cells/μm$^2$ | 0.0049 | ** |
| | | | | | | | CA1 (Young) vs. CA2 (Aged) | 0.0000944 | 0.000008325 | 0.0002832 | 0.00003419 | # of cells/μm$^2$ | <0.0001 | **** |
| | | | | | | | CA1 (Young) vs. CA3 (Young) | 0.0000944 | 0.000008325 | 0.0001626 | 0.0000346 | # of cells/μm$^2$ | 0.4268 | ns |
| | | | | | | | CA1 (Young) vs. CA3 (Aged) | 0.0000944 | 0.000008325 | 0.000183 | 0.0000324 | # of cells/μm$^2$ | 0.1318 | ns |
| | | | | | | | CA1 (Young) vs. DG (Young) | 0.0000944 | 0.000008325 | 0.0001194 | 0.00001358 | # of cells/μm$^2$ | 0.9944 | ns |
| | | | | | | | CA1 (Young) vs. DG (Aged) | 0.0000944 | 0.000008325 | 0.0001271 | 0.00001779 | # of cells/μm$^2$ | 0.9737 | ns |
| | | | | | | | CA1 (Aged) vs. CA2 (Young) | 0.0001058 | 0.00001013 | 0.0002196 | 0.00002244 | # of cells/μm$^2$ | 0.0161 | * |
| | | | | | | | CA1 (Aged) vs. CA2 (Aged) | 0.0001058 | 0.00001013 | 0.0002832 | 0.00003419 | # of cells/μm$^2$ | <0.0001 | **** |
| | | | | | | | CA1 (Aged) vs. CA3 (Young) | 0.0001058 | 0.00001013 | 0.0001626 | 0.0000346 | # of cells/μm$^2$ | 0.6645 | ns |
| | | | | | | | CA1 (Aged) vs. CA3 (Aged) | 0.0001058 | 0.00001013 | 0.000183 | 0.0000324 | # of cells/μm$^2$ | 0.2777 | ns |
| | | | | | | | CA1 (Aged) vs. DG (Young) | 0.0001058 | 0.00001013 | 0.0001194 | 0.00001358 | # of cells/μm$^2$ | 0.9999 | ns |
| | | | | | | | CA1 (Aged) vs. DG (Aged) | 0.0001058 | 0.00001013 | 0.0001271 | 0.00001779 | # of cells/μm$^2$ | 0.9981 | ns |
| | | | | | | | CA2 (Young) vs. CA2 (Aged) | 0.0002196 | 0.00002244 | 0.0002832 | 0.00003419 | # of cells/μm$^2$ | 0.5255 | ns |
| | | | | | | | CA2 (Young) vs. CA3 (Young) | 0.0002196 | 0.00002244 | 0.0001626 | 0.0000346 | # of cells/μm$^2$ | 0.6567 | ns |
| | | | | | | | CA2 (Young) vs. CA3 (Aged) | 0.0002196 | 0.00002244 | 0.000183 | 0.0000324 | # of cells/μm$^2$ | 0.9517 | ns |
| | | | | | | | CA2 (Young) vs. DG (Young) | 0.0002196 | 0.00002244 | 0.0001194 | 0.00001358 | # of cells/μm$^2$ | 0.0526 | ns |
| | | | | | | | CA2 (Young) vs. DG (Aged) | 0.0002196 | 0.00002244 | 0.0001271 | 0.00001779 | # of cells/μm$^2$ | 0.0995 | ns |
| | | | | | | | CA2 (Aged) vs. CA3 (Young) | 0.0002832 | 0.00003419 | 0.0001626 | 0.0000346 | # of cells/μm$^2$ | 0.0083 | ** |
| | | | | | | | CA2 (Aged) vs. CA3 (Aged) | 0.0002832 | 0.00003419 | 0.000183 | 0.0000324 | # of cells/μm$^2$ | 0.056 | ns |
| | | | | | | | CA2 (Aged) vs. DG (Young) | 0.0002832 | 0.00003419 | 0.0001194 | 0.00001358 | # of cells/μm$^2$ | <0.0001 | **** |
| | | | | | | | CA2 (Aged) vs. DG (Aged) | 0.0002832 | 0.00003419 | 0.0001271 | 0.00001779 | # of cells/μm$^2$ | 0.0002 | *** |
| | | | | | | | CA3 (Young) vs. CA3 (Aged) | 0.0001626 | 0.0000346 | 0.000183 | 0.0000324 | # of cells/μm$^2$ | 0.9985 | ns |
| | | | | | | | CA3 (Young) vs. DG (Young) | 0.0001626 | 0.0000346 | 0.0001194 | 0.00001358 | # of cells/μm$^2$ | 0.8879 | ns |
| | | | | | | | CA3 (Young) vs. DG (Aged) | 0.0001626 | 0.0000346 | 0.0001271 | 0.00001779 | # of cells/μm$^2$ | 0.9589 | ns |
| | | | | | | | CA3 (Aged) vs. DG (Young) | 0.000183 | 0.0000324 | 0.0001194 | 0.00001358 | # of cells/μm$^2$ | 0.5263 | ns |
| | | | | | | | CA3 (Aged) vs. DG (Aged) | 0.000183 | 0.0000324 | 0.0001271 | 0.00001779 | # of cells/μm$^2$ | 0.6876 | ns |
| | | | | | | | DG (Young) vs. DG (Aged) | 0.0001194 | 0.00001358 | 0.0001271 | 0.00001779 | # of cells/μm$^2$ | >0.9999 | ns |
| **Figure 3** | **Subfigure** | **Experiment** | **Strain** | **Young** | **Aged** | **Statistical test** | **Comparison** | **Young Mean** | **SEM** | **Aged Mean** | **SEM** | **Units** | **p-value** | **Significance** |
| 3 | B | Dorsal striatum cell counts | C57/BL6 | N=5, 5-8 sections/ mouse | N=5, 4-6 sections/ mouse | Student t-test | PV$^+$ Cells | 36.97 | 4.323 | 37.83 | 2.641 | # of cells/mm$^2$ | 0.8697 | ns |
| | | | | | | | WFA$^+$ Cells | 23.94 | 4.733 | 28.3 | 1.46 | # of cells/mm$^2$ | 0.4048 | ns |
| | | | | | | | ACAN$^+$ Cells | 9.538 | 2.039 | 6.139 | 0.3701 | # of cells/mm$^2$ | 0.1071 | ns |
| | | | | | | | PV$^+$:WFA$^+$ Cells | 16.14 | 3.782 | 18.48 | 1.35 | # of cells/mm$^2$ | 0.5762 | ns |
| | | | | | | | PV$^+$:ACAN$^+$ Cells | 1.009 | 0.1641 | 0.9083 | 0.1583 | # of cells/mm$^2$ | 0.6693 | ns |
| | | | | | | | WFA$^+$:ACAN$^+$ Cells | 0.691 | 0.1275 | 0.6805 | 0.05578 | # of cells/mm$^2$ | 0.9419 | ns |
| | | | | | | | PV$^+$:WFA$^+$:ACAN$^+$ Cells | 0.532 | 0.1155 | 0.4588 | 0.06843 | # of cells/mm$^2$ | 0.6009 | ns |
| | C | Percentage of PV$^+$ Cells with a WFA$^+$ PNN (Total striatum) | C57/BL6 | N=5, 5-8 sections/ mouse | N=5, 4-6 sections/ mouse | Student t-test | Young vs Aged | 40.8 | 4.118 | 49.15 | 1.271 | % of PV cells | 0.0886 | ns |
| | | | | | | | **[A] vs [B]** | **[A] Mean** | **[A] SEM** | **[B] Mean** | **[B] SEM** | **Units** | **p-value** | **Significance** |
| | D | PV+ Cells with a WFA+ PNN (Striatal regions) | C57/BL6 | N=5, 5-8 sections/ mouse | N=5, 4-6 sections/ mouse | Two-way ANOVA w/ Tukey | DMS (Young) vs DLS (Young) | 22.07 | 1.986 | 39.44 | 4.552 | % of PV cells | <0.0001 | **** |
| | | | | | | | DMS (Young) vs DMS (Aged) | 22.07 | 1.986 | 21.7 | 2.037 | % of PV cells | 0.9992 | ns |
| | | | | | | | DMS (Young) vs DLS (Aged) | 22.07 | 1.986 | 27.41 | 2.306 | % of PV cells | 0.2568 | ns |
| | | | | | | | DLS (Young) vs DMS (Aged) | 39.44 | 4.552 | 21.7 | 2.037 | % of PV cells | <0.0001 | **** |
| | | | | | | | DLS (Young) vs DLS (Aged) | 39.44 | 4.552 | 27.41 | 2.306 | % of PV cells | 0.0005 | *** |
| | | | | | | | DMS (Aged) vs DLS (Aged) | 21.7 | 2.037 | 27.41 | 2.306 | % of PV cells | 0.1529 | ns |

| Figure 3 | Subfigure | Experiment | Strain | Young | Aged | Statistical test | Comparison | Young Mean | SEM | Aged Mean | SEM | Units | p-value | Significance |
|---|---|---|---|---|---|---|---|---|---|---|---|---|---|---|
| 3 | | | | | | | [A] vs [B] | [A] Mean | [A] SEM | [B] Mean | [B] SEM | Units | p-value | Significance |
| | E | Percentage of PV+ Cells with a WFA+ PNN (Striatal regions) | C57/BL6 | N=5, 5-8 sections/ mouse | N=5, 4-6 sections/ mouse | Two-way ANOVA w/ Tukey | DMS (Young) vs DLS (Young) | 38.19 | 3.572 | 44.34 | 4.776 | % of PV cells | 0.2719 | ns |
| | | | | | | | DMS (Young) vs DMS (Aged) | 38.19 | 3.572 | 50.4 | 1.777 | % of PV cells | 0.0058 | ** |
| | | | | | | | DMS (Young) vs DLS (Aged) | 38.19 | 3.572 | 48.67 | 0.9025 | % of PV cells | 0.0239 | * |
| | | | | | | | DLS (Young) vs DMS (Aged) | 44.34 | 4.776 | 50.4 | 1.777 | % of PV cells | 0.3421 | ns |
| | | | | | | | DLS (Young) vs DLS (Aged) | 44.34 | 4.776 | 48.67 | 0.9025 | % of PV cells | 0.6308 | ns |
| | | | | | | | DMS (Aged) vs DLS (Aged) | 50.4 | 1.777 | 48.67 | 0.9025 | % of PV cells | 0.9688 | ns |
| | F | Colocalization of $PV^+$:$WFA^+$ | C57/BL6 | N=5, 2 sections/ mouse | N=5, 2 sections/ mouse | Two-way ANOVA w/ Tukey | DMS (Young) vs DLS (Young) | 0.2629 | 0.2915 | 0.4233 | 0.02819 | Pearson's r | 0.0011 | ** |
| | | | | | | | DMS (Young) vs DMS (Aged) | 0.2629 | 0.2915 | 0.366 | 0.02362 | Pearson's r | 0.053 | ns |
| | | | | | | | DMS (Young) vs DLS (Aged) | 0.2629 | 0.2915 | 0.4798 | 0.02539 | Pearson's r | <0.0001 | **** |
| | | | | | | | DLS (Young) vs DMS (Aged) | 0.4233 | 0.2819 | 0.366 | 0.02362 | Pearson's r | 0.4112 | ns |
| | | | | | | | DLS (Young) vs DLS (Aged) | 0.4233 | 0.2819 | 0.4798 | 0.02539 | Pearson's r | 0.4076 | ns |
| | | | | | | | DMS (Aged) vs DLS (Aged) | 0.366 | 0.02362 | 0.4798 | 0.02539 | Pearson's r | 0.0256 | * |
| Figure 4 | Subfigure | Experiment | Region | Young | Aged | Statistical test | Age group | Equation | | $R^2$ | Y-intercept | Slope | p-value | Significance |
| 4 | A | Linear regression of $WFA^+$ vs $PV^+$ intensity | Striatum | N=5, 2-7 sections/ mouse | N=5, 3-8 sections/ mouse | Linear regression/ Coefficient difference t-test of slopes | Young | Y = 0.7525*X +0.2125 | | 0.5681 | 0.2125 | 0.7525 | 0.1412 | ns |
| | | | | | | | Aged | Y = 0.9409*X + 0.08875 | | 0.974 | 0.08875 | 0.9409 | 0.0018 | ** |
| | B | | Hippocampus | | | | Young | Y = 1.011*X + 0.01704 | | 0.3062 | 0.01704 | 1.011 | 0.0093 | ** |
| | | | | | | | Aged | Y = 0.7325*X + 0.2406 | | 0.4345 | 0.2406 | 0.7325 | 0.0029 | ** |
| | C | Linear regression of $WFA^+$ vs $ACAN^+$ intensity | Striatum | | | | Young | Y = 0.001760*X + 0.7824 | | 0.000002881 | 0.7824 | 0.00176 | 0.9978 | ns |
| | | | | | | | Aged | Y = 0.4923*X + 0.4269 | | 0.5271 | 0.4269 | 0.4923 | 0.1649 | ns |
| | D | | Hippocampus | | | | Young | Y = 0.09677*X + 0.7287 | | 0.005889 | 0.7287 | 0.09677 | 0.7409 | ns |
| | | | | | | | Aged | Y = 0.5204*X + 0.4462 | | 0.5328 | 0.4462 | 0.5204 | 0.0006 | *** |
| Figure 5 (See supplementary table 2) | | | | | | | | | | | | | | |
| Figure 6 | Subfigure | Experiment | Strain | Young | Aged | Statistical test | [A] vs [B] | [A] Mean | [A] SEM | [B] Mean | [B] SEM | Units | p-value | Significance |
| 6 | B | Iba1 Intensity per cell | C57/BL6 | N=5, 4-17 cells/ region/ mouse | N=5, 4-15 cells/ region/ mouse | Two-way ANOVA w/ Tukey | Str. (Young) vs Str. (Aged) | 42.07 | 3.766 | 64.66 | 7.634 | A.U./cell | <0.0001 | **** |
| | | | | | | | Str. (Young) vs Hpc. (Young) | 42.07 | 3.766 | 32.56 | 4.155 | A.U./cell | 0.3364 | ns |
| | | | | | | | Str. (Young) vs Hpc. (Aged) | 42.07 | 3.766 | 47.45 | 5.76 | A.U./cell | 0.7456 | ns |
| | | | | | | | Str. (Aged) vs Hpc. (Young) | 64.66 | 7.634 | 32.56 | 4.155 | A.U./cell | <0.0001 | **** |
| | | | | | | | Str. (Aged) vs Hpc. (Aged) | 64.66 | 7.634 | 47.45 | 5.76 | A.U./cell | 0.0089 | ** |
| | | | | | | | Str. (Young) vs Hpc. (Young) | 32.56 | 4.155 | 47.45 | 5.76 | A.U./cell | 0.0959 | ns |
| | C | Striatal mean Iba1 intensity | C57/BL6 | N=5, 3-4 sections/ mouse | N=5, 4 sections/ mouse | Student t-test | Young vs Aged | 202.9 | 3.82 | 215.3 | 3.245 | A.U. | 0.0391 | * |
| | D | Striatal Iba1 intensity per cell | C57/BL6 | N=5, 2-12 cells/ region/ mouse | N=5, 3-12 cells/ region/ mouse | Two-way ANOVA w/ Tukey | DMS (Young) vs DLS (Young) | 47.05 | 8.569 | 40.47 | 4.353 | A.U./cell | 0.6523 | ns |
| | | | | | | | DMS (Young) vs DMS (Aged) | 47.05 | 8.569 | 54.74 | 7.97 | A.U./cell | 0.557 | ns |
| | | | | | | | DMS (Young) vs DLS (Aged) | 47.05 | 8.569 | 72.21 | 12.19 | A.U./cell | 0.0005 | *** |
| | | | | | | | DLS (Young) vs DMS (Aged) | 40.47 | 4.353 | 54.74 | 7.97 | A.U./cell | 0.032 | * |
| | | | | | | | DLS (Young) vs DLS (Aged) | 40.47 | 4.353 | 72.21 | 12.19 | A.U./cell | <0.0001 | **** |
| | | | | | | | DMS (Aged) vs DLS (Aged) | 54.74 | 7.97 | 72.21 | 12.19 | A.U./cell | 0.0128 | * |
| | E | Average cell body area | C57/BL6 | N=5, 4-17 cells/ region/ mouse | N=5, 4-15 cells/ region/ mouse | Two-way ANOVA w/ Tukey | Str. (Young) vs Str. (Aged) | 36.36 | 2.217 | 51.73 | 3.09 | $\mu m^2$ | <0.0001 | **** |
| | | | | | | | Str. (Young) vs Hpc. (Young) | 36.36 | 2.217 | 37.07 | 2.388 | $\mu m^2$ | 0.9978 | ns |
| | | | | | | | Str. (Young) vs Hpc. (Aged) | 36.36 | 2.217 | 40.5 | 4.369 | $\mu m^2$ | 0.6817 | ns |
| | | | | | | | Str. (Aged) vs Hpc. (Young) | 51.73 | 3.09 | 37.07 | 2.388 | $\mu m^2$ | 0.0022 | ** |
| | | | | | | | Str. (Aged) vs Hpc. (Aged) | 51.73 | 3.09 | 40.5 | 4.369 | $\mu m^2$ | 0.0227 | * |
| | | | | | | | Hpc. (Young) vs Hpc. (Aged) | 37.07 | 2.388 | 40.5 | 4.369 | $\mu m^2$ | 0.8782 | ns |
| | F | Average cell body perimeter | C57/BL6 | N=5, 4-17 cells/ region/ mouse | N=5, 4-15 cells/ region/ mouse | Two-way ANOVA w/ Tukey | Str. (Young) vs Str. (Aged) | 23.72 | 0.7215 | 29.45 | 1.1 | μm | <0.0001 | **** |
| | | | | | | | Str. (Young) vs Hpc. (Young) | 23.72 | 0.7215 | 24.22 | 0.9237 | μm | 0.9871 | ns |
| | | | | | | | Str. (Young) vs Hpc. (Aged) | 23.72 | 0.7215 | 26.51 | 1.971 | μm | 0.2028 | ns |
| | | | | | | | Str. (Aged) vs Hpc. (Young) | 29.45 | 1.1 | 24.22 | 0.9237 | μm | 0.0035 | ** |
| | | | | | | | Str. (Aged) vs Hpc. (Aged) | 29.45 | 1.1 | 26.51 | 1.971 | μm | 0.1721 | ns |
| | | | | | | | Str. (Young) vs Hpc. (Young) | 24.22 | 0.9237 | 26.51 | 1.971 | μm | 0.5338 | ns |
| | G | Striatal cell body area | C57/BL6 | N=5, 2-12 cells/ region/ mouse | N=5, 3-12 cells/ region/ mouse | Two-way ANOVA w/ Tukey | DMS (Young) vs DLS (Young) | 36.47 | 2.478 | 38.12 | 2.034 | $\mu m^2$ | 0.9849 | ns |
| | | | | | | | DMS (Young) vs DMS (Aged) | 36.47 | 2.478 | 48.15 | 2.874 | $\mu m^2$ | 0.0674 | ns |
| | | | | | | | DMS (Young) vs DLS (Aged) | 36.47 | 2.478 | 57.03 | 5.583 | $\mu m^2$ | 0.0005 | *** |
| | | | | | | | DLS (Young) vs DMS (Aged) | 38.12 | 2.034 | 48.15 | 2.874 | $\mu m^2$ | 0.0693 | ns |
| | | | | | | | DLS (Young) vs DLS (Aged) | 38.12 | 2.034 | 57.03 | 5.583 | $\mu m^2$ | 0.0003 | *** |
| | | | | | | | DMS (Aged) vs DLS (Aged) | 48.15 | 2.874 | 57.03 | 5.583 | $\mu m^2$ | 0.1997 | ns |
| | H | Striatal cell body perimeter | C57/BL6 | N=5, 2-12 cells/ region/ mouse | N=5, 3-12 cells/ region/ mouse | Two-way ANOVA w/ Tukey | DMS (Young) vs DLS (Young) | 23.55 | 0.9868 | 24.2 | 0.7286 | μm | 0.9813 | ns |
| | | | | | | | DMS (Young) vs DMS (Aged) | 23.55 | 0.9868 | 28.71 | 1.256 | μm | 0.0216 | * |
| | | | | | | | DMS (Young) vs DLS (Aged) | 23.55 | 0.9868 | 30.54 | 1.321 | μm | 0.0014 | ** |
| | | | | | | | DLS (Young) vs DMS (Aged) | 24.2 | 0.7286 | 28.71 | 1.256 | μm | 0.0219 | * |
| | | | | | | | DLS (Young) vs DLS (Aged) | 24.2 | 0.7286 | 30.54 | 1.321 | μm | 0.001 | ** |
| | | | | | | | DMS (Aged) vs DLS (Aged) | 28.71 | 1.256 | 30.54 | 1.321 | μm | 0.7006 | ns |
| | | | | | | | Group | Branch | | No Branch | | Units | | |
| | I | Microglia morphology characterization | C57/BL6 | N=5, 25-64 cells/mouse | N=5, 29-58 cells/mouse | No statistical tests | Striatum (Young) | 30 | 46.88% | 34 | 53.13% | # of cells | N.A. | |
| | | | | | | | Striatum (Aged) | 19 | 32.76% | 39 | 67.24% | # of cells | | |
| | | | | | | | Hippocampus (Young) | 20 | 80.00% | 5 | 20.00% | # of cells | | |
| | | | | | | | Hippocampus (Aged) | 20 | 68.97% | 9 | 31.03% | # of cells | | |
| | | | | | | | [A] vs [B] | [A] Mean | [A] SEM | [B] Mean | [B] SEM | Units | p-value | Significance |
| | J | Average number of endpoints | C57/BL6 | N=5, 4-17 cells/ region/ mouse | N=5, 4-15 cells/ region/ mouse | Two-way ANOVA w/ Tukey | Str. (Young) vs Str. (Aged) | 8.997 | 0.634 | 6.951 | 0.9349 | # of endpoints | * | 0.0293 |
| | | | | | | | Str. (Young) vs Hpc. (Young) | 8.997 | 0.634 | 5.193 | 0.7651 | # of endpoints | *** | 0.0005 |
| | | | | | | | Str. (Young) vs Hpc. (Aged) | 8.997 | 0.634 | 4.406 | 0.641 | # of endpoints | **** | <0.0001 |
| | | | | | | | Str. (Aged) vs Hpc. (Young) | 6.951 | 0.9349 | 5.193 | 0.7651 | # of endpoints | ns | 0.2809 |
| | | | | | | | Str. (Aged) vs Hpc. (Aged) | 6.951 | 0.9349 | 4.406 | 0.641 | # of endpoints | * | 0.0384 |
| | | | | | | | Str. (Young) vs Hpc. (Young) | 5.193 | 0.7651 | 4.406 | 0.641 | # of endpoints | ns | 0.8946 |
| | K | Sholl analysis | C57/BL6 | N=5, 5-26 cells/ mouse | N=5, 5-29 cells/ mouse | Three-way ANOVA w/ Tukey | Source of Variation | SS (Type III) | DF | MS | F (DFn, DFd) | | p-value | Significance |
| | | | | | | | Distance from Soma | 367.8 | 19 | 19.36 | F (19, 1220) = 39.62 | | <0.0001 | **** |
| | | | | | | | Region | 7.732 | 1 | 7.732 | F (1, 1220) = 15.82 | | <0.0001 | **** |
| | | | | | | | Age | 0.4463 | 1 | 0.4463 | F (1, 1220) = 0.9134 | | 0.3394 | ns |
| | | | | | | | Distance from Soma x Region | 72.53 | 19 | 3.817 | F (19, 1220) = 7.812 | | <0.0001 | **** |
| | | | | | | | Distance from Soma x Age | 2.543 | 19 | 0.1338 | F (19, 1220) = 0.2739 | | 0.9992 | ns |
| | | | | | | | Region x Age | 0.2107 | 1 | 0.2107 | F (1, 1220) = 0.4313 | | 0.5115 | ns |
| | | | | | | | Distance from Soma x Region x A | 1.9 | 19 | 0.1 | F (19, 1220) = 0.2047 | | >0.9999 | ns |
| | | | | | | Three-way ANOVA w/ Tukey by region | Distance from Soma (μm) | Young-Str | Young-Hpc | Aged-Str | Aged-Hpc | p-value (Hpc.) | p-value (Str.) | Significance |
| | | | | | | | 0 | 0 | 1.8 | 0 | 1.655172414 | 0.4702 | >0.9999 | ns |
| | | | | | | | 5 | 3.765757576 | 2.32 | 3.225826649 | 1.965517241 | 0.0773 | 0.0022 | ## |
| | | | | | | | 10 | 2.943030303 | 1.12 | 2.164455214 | 1.206896552 | 0.6648 | <0.0001 | #### |
| | | | | | | | 15 | 1.706969697 | 0.56 | 1.926363636 | 0.517241379 | 0.8311 | 0.2088 | ns |
| | | | | | | | 20 | 1.07969697 | 0.16 | 0.992942291 | 0.275862069 | 0.5634 | 0.6184 | ns |
| | | | | | | | 25 | 0.78969697 | 0.12 | 0.558786765 | 0.103448276 | 0.9342 | 0.186 | ns |
| | | | | | | | 30 | 0.35 | 0 | 0.223902629 | 0.034482759 | 0.8635 | 0.4693 | ns |
| | | | | | | | 35 | 0.206666667 | 0 | 0.164622326 | 0 | >0.9999 | 0.8092 | ns |
| | | | | | | | 40 | 0.166666667 | 0 | 0.071657754 | 0 | >0.9999 | 0.5855 | ns |
| | | | | | | | 45 | 0.04 | 0 | 0 | 0 | >0.9999 | 0.8183 | ns |
| | | | | | | | 50 | 0.013333333 | 0 | 0 | 0 | >0.9999 | 0.939 | ns |
| | | | | | | | 55-100 | 0 | 0 | 0 | 0 | >0.9999 | >0.9999 | ns |

### Supplementary Table 2

| | | | | | | | | Hippocampus | | | | | | Striatum | | | | | |
|---|---|---|---|---|---|---|---|---|---|---|---|---|---|---|---|---|---|---|---|
| | | | | | | | | Young | | Aged | | p-value | Significance | Young | | Aged | | p-value | Significance |
| Figure 5 | Strain | Young | Aged | Statistical test | Subfigure | Experiment | Marker | Fold Change | ΔCt | Fold Change | ΔCt | | | Fold Change | ΔCt | Fold Change | ΔCt | | |
| 5 | C57/BL6 | N=5 | N=5 | Student t-test | A | Glial marker expression | *GFAP* | 1.064 | 14.43 | 1.814 | 13.64 | 0.0514 | ns | 1.018 | 10.57 | 2.428 | 9.382 | 0.0031 | ** |
| | | | | | | | *AQP4* | 1.015 | 16.42 | 1.003 | 16.44 | 0.9120 | ns | 1.004 | 11.17 | 1.176 | 10.97 | 0.2289 | ns |
| | | | | | | | *Iba1* | 1.030 | 18.87 | 1.307 | 18.51 | 0.1278 | ns | 1.007 | 13.13 | 1.480 | 12.59 | 0.0075 | ** |
| | | | | | | | *TREM2* | 1.036 | 17.99 | 1.251 | 17.68 | 0.1855 | ns | 1.003 | 12.68 | 1.524 | 12.12 | 0.0206 | * |
| | | | | | | | *CD68* | 1.016 | 11.19 | 1.159 | 10.98 | 0.1472 | ns | 1.006 | 10.79 | 1.407 | 10.31 | 0.0026 | ** |
| | | | | | | | *TLR2* | 1.015 | 16.68 | 2.209 | 15.61 | 0.0033 | ** | 1.015 | 16.68 | 2.209 | 15.61 | 0.0033 | ** |
| | | | | | | | *TLR4* | 1.013 | 16.24 | 1.291 | 15.89 | 0.0508 | ns | 1.018 | 16.21 | 1.266 | 15.89 | 0.1001 | ns |
| | | | | | B | Neuroinflammatory marker expression | *IL-1β* | 1.024 | 24.48 | 2.335 | 23.33 | 0.0046 | ** | 1.047 | 19.83 | 2.022 | 19.05 | 0.1150 | ns |
| | | | | | | | *IL-6* | 1.027 | 18.05 | 1.761 | 17.24 | 0.0022 | ** | 1.021 | 18.23 | 2.133 | 17.14 | 0.0002 | *** |
| | | | | | | | *IL-10* | Failed assumptions of ΔΔCt Method | | | | NA | ND | Failed assumptions of ΔΔCt Method | | | | NA | ND |
| | | | | | | | *TNF-α* | 1.090 | 24.97 | 2.923 | 23.50 | 0.0056 | ** | 1.066 | 18.99 | 3.707 | 17.28 | 0.0054 | ** |
| | | | | | | | *TGF-β* | 1.021 | 20.43 | 1.219 | 20.15 | 0.1062 | ns | 1.009 | 20.05 | 1.073 | 19.95 | 0.4233 | ns |
| | | | | | | | *CCL2* | 1.055 | 24.45 | 2.714 | 23.04 | 0.0013 | ** | 1.169 | 19.17 | 2.950 | 17.76 | 0.0286 | * |
| | | | | | | | *CCL5* | 1.007 | 16.71 | 1.774 | 16.04 | 0.0756 | ns | 1.027 | 16.54 | 2.508 | 15.36 | 0.0126 | * |
| | | | | | C | Complement and senescence-related marker expression | *C1qa* | 1.007 | 9.029 | 1.251 | 8.569 | 0.0085 | ** | 1.000 | 8.871 | 1.353 | 8.445 | 0.0012 | ** |
| | | | | | | | *C3* | 1.019 | 14.76 | 2.595 | 13.58 | 0.0163 | * | 1.090 | 15.60 | 4.291 | 13.62 | 0.0015 | ** |
| | | | | | | | *C3ar1* | 1.007 | 14.92 | 1.383 | 14.60 | 0.0253 | * | 1.006 | 14.76 | 1.360 | 14.36 | 0.0912 | ns |
| | | | | | | | *P16* | 1.027 | 24.85 | 2.874 | 23.37 | 0.0002 | *** | 1.052 | 13.99 | 2.737 | 12.55 | 0.0004 | *** |
| | | | | | | | *P21* | 1.003 | 13.08 | 1.045 | 13.03 | 0.6630 | ns | 1.003 | 13.09 | 0.981 | 13.14 | 0.7251 | ns |
| | | | | | D | PNN component and matrix metalloproteinase marker expression | *HAPLN1* | 1.024 | 18.51 | 1.044 | 18.47 | 0.8103 | ns | 1.006 | 22.86 | 1.001 | 22.89 | 0.8450 | ns |
| | | | | | | | *PV* | 1.069 | 15.44 | 0.880 | 15.66 | 0.4806 | ns | 1.020 | 10.29 | 1.112 | 10.16 | 0.5205 | ns |
| | | | | | | | *ACAN* | 1.014 | 20.77 | 1.094 | 20.66 | 0.5009 | ns | 1.014 | 15.70 | 1.236 | 15.42 | 0.1167 | ns |
| | | | | | | | *MMP3* | 0.836 | 19.94 | 18.340 | 15.25 | 0.0004 | *** | 1.259 | 18.15 | 2.336 | 17.60 | 0.5605 | ns |
| | | | | | | | *MMP3 (Retest)* | 1.085 | 16.35 | 19.341 | 12.24 | <0.0001 | **** | 1.441 | 15.69 | 3.001 | 15.20 | 0.6241 | ns |
| | | | | | | | *MMP9* | 1.007 | 14.80 | 0.844 | 15.06 | 0.0835 | ns | 1.009 | 16.14 | 1.274 | 15.80 | 0.0461 | * |
| | | | | | | | *TIMP1* | 1.021 | 17.62 | 1.749 | 16.84 | 0.0062 | ** | 1.129 | 17.74 | 1.811 | 17.13 | 0.2841 | ns |
| | | | | | | | *ADAM17* | 1.003 | 8.397 | 0.8422 | 8.65 | 0.0165 | * | 1.001 | 10.72 | 1.205 | 10.46 | 0.0222 | * |
| | | | | | Not shown | Other markers tested but not shown in Figure 5 | *IFN-γ* | 1.047 | 18.92 | 1.000 | 19.72 | 0.4438 | ns | 1.017 | 18.61 | 0.994 | 19.01 | 0.5365 | ns |
| | | | | | | | *TMEM119* | 1.004 | 12.17 | 1.211 | 11.92 | 0.0980 | ns | 1.004 | 12.17 | 1.211 | 11.92 | 0.0980 | ns |
| | | | | | | | *MMP2* | 1.050 | 15.07 | 0.869 | 15.28 | 0.3763 | ns | 1.013 | 15.42 | 1.140 | 15.25 | 0.3254 | ns |
| | | | | | | | *MMP13* | 1.015 | 17.18 | 0.674 | 18.00 | 0.0971 | ns | 1.418 | 17.26 | 0.654 | 18.02 | 0.2661 | ns |
| | | | | | | | *TLR3* | 1.008 | 13.17 | 1.121 | 13.02 | 0.3156 | ns | 1.009 | 13.18 | 1.130 | 13.02 | 0.2870 | ns |
| | | | | | | | *TLR5* | 1.032 | 17.34 | 1.160 | 17.17 | 0.5130 | ns | 1.047 | 17.30 | 1.175 | 17.12 | 0.5490 | ns |
| | | | | | | | *NFE212* | 1.005 | 13.23 | 1.171 | 13.02 | 0.1382 | ns | 1.005 | 13.23 | 1.171 | 13.02 | 0.1328 | ns |

| Supplementary Table 3 | | | | | |
|---|---|---|---|---|---|
| **Gene** | **Gene name** | **Company** | **Cat. #** | **Species** | **Color** |
| ***Acan*** | ***Aggrecan*** | Applied Biosystems-ThermoFisher | Mm00645794_m1 | Mouse | FAM |
| ***Adam17*** | ***A disintegrin and metallopeptidase domain 17*** | Applied Biosystems-ThermoFisher | Mm00456428_m1 | Mouse | FAM |
| ***Aif (Iba1)*** | ***Allograt inflammatory factor 1/Ionized calcium-binding adaptor molecule 1*** | Applied Biosystems-ThermoFisher | Mm00479862_g1 | Mouse | FAM |
| ***Aqp4*** | ***Aquaporin 4*** | Applied Biosystems-ThermoFisher | Mm00802131_m1 | Mouse | FAM |
| ***C1qa*** | ***Complement component 1, q subcomponen, alpha polypeptide*** | Applied Biosystems-ThermoFisher | Mm00432142_m1 | Mouse | FAM |
| ***C3*** | ***Complement component 3*** | Applied Biosystems-ThermoFisher | Mm01232779_m1 | Mouse | FAM |
| ***C3ar1*** | ***Complement component 3a receptor 1*** | Applied Biosystems-ThermoFisher | Mm02620006_s1 | Mouse | FAM |
| ***Ccl2*** | ***Chemokine (C-C motif) ligand 2*** | Applied Biosystems-ThermoFisher | Mm00441242_m1 | Mouse | FAM |
| ***Ccl5*** | ***Chemokine (C-C motif) ligand 5*** | Applied Biosystems-ThermoFisher | Mm01302427_m1 | Mouse | FAM |
| ***Cd68*** | ***CD68 antigen*** | Applied Biosystems-ThermoFisher | Mm03047343_m1 | Mouse | FAM |
| ***Cdkn1a (P21)*** | ***Cyclin-dependent kinase inhibitor 1A*** | Applied Biosystems-ThermoFisher | Mm04205640_g1 | Mouse | FAM |
| ***Cdkn2a (P16)*** | ***Cyclin-dependent kinase inhibitor 2A*** | Applied Biosystems-ThermoFisher | Mm00494449_m1 | Mouse | FAM |
| ***Gapdh*** | ***Glyceraldehyde-3-phosphate dehydrogenase*** | Applied Biosystems-ThermoFisher | Mm99999915_g1 | Mouse | FAM |
| ***Gfap*** | ***Glial fibrillary acidic protein*** | Applied Biosystems-ThermoFisher | Mm01253030_m1 | Mouse | FAM |
| ***Hapln1*** | ***Hyaluronan and proteoglycan link protein 1*** | Applied Biosystems-ThermoFisher | Mm00488952_m1 | Mouse | FAM |
| ***Ifng*** | ***Interferon gamma*** | Applied Biosystems-ThermoFisher | Mm01168134_m1 | Mouse | FAM |
| ***Il1b*** | ***Interleukin 1 beta*** | Applied Biosystems-ThermoFisher | Mm00434228_m1 | Mouse | FAM |
| ***Il2*** | ***Interleukin 2*** | Applied Biosystems-ThermoFisher | Mm00434256_m1 | Mouse | FAM |
| ***Il6*** | ***Interleukin 6*** | Applied Biosystems-ThermoFisher | Mm00446190_m1 | Mouse | FAM |
| ***Il10*** | ***Interleukin 10*** | Applied Biosystems-ThermoFisher | Mm00439614_m1 | Mouse | FAM |
| ***Mmp2*** | ***Matrix metallopeptidase 2*** | Applied Biosystems-ThermoFisher | Mm00439498_m1 | Mouse | FAM |
| ***Mmp3*** | ***Matrix metallopeptidase 3*** | Applied Biosystems-ThermoFisher | Mm00440295_m1 | Mouse | FAM |
| ***Mmp9*** | ***Matrix metallopeptidase 9*** | Applied Biosystems-ThermoFisher | Mm00442991_m1 | Mouse | FAM |
| ***Mmp13*** | ***Matrix metallopeptidase 13*** | Applied Biosystems-ThermoFisher | Mm00439491_m1 | Mouse | FAM |
| ***Nfe2l2*** | ***Nuclear factor, erythroid derived 2, like 2*** | Applied Biosystems-ThermoFisher | Mm00477784_m1 | Mouse | FAM |
| ***Pvalb (PV)*** | ***Parvalbumin*** | Applied Biosystems-ThermoFisher | Mm0043100_m1 | Mouse | FAM |
| ***Timp1*** | ***Tissue inhibitor of metalloproteinase 1*** | Applied Biosystems-ThermoFisher | Mm01341361_m1 | Mouse | FAM |
| ***Tgfb1*** | ***Transforming growth factor, beta 1*** | Applied Biosystems-ThermoFisher | Mm01178820_m1 | Mouse | FAM |
| ***Tmem119*** | ***Transmembrane protein 119*** | Applied Biosystems-ThermoFisher | Mm00525305_m1 | Mouse | FAM |
| ***Tnf*** | ***Tumor necrosis facter alpha*** | Applied Biosystems-ThermoFisher | Mm00443258_m1 | Mouse | FAM |
| ***Tlr2*** | ***Toll-like receptor 2*** | Applied Biosystems-ThermoFisher | Mm00442346_m1 | Mouse | FAM |
| ***Tlr3*** | ***Toll-like receptor 3*** | Applied Biosystems-ThermoFisher | Mm01207404_m1 | Mouse | FAM |
| ***Tlr4*** | ***Toll-like receptor 4*** | Applied Biosystems-ThermoFisher | Mm00445273_m1 | Mouse | FAM |
| ***Tlr5*** | ***Toll-like receptor 5*** | Applied Biosystems-ThermoFisher | Mm00546288_s1 | Mouse | FAM |
| ***Trem2*** | ***Triggering receptor expressed on myeloid cells 2*** | Applied Biosystems-ThermoFisher | Mm04209422_m1 | Mouse | FAM |
| ***18S*** | ***Eukaryotic 18S rRNA*** | Applied Biosystems-ThermoFisher | 4333760F | Eukaryotic | FAM |

### Supplementary Table 4

| Figure(s) | Experiment | Microscope Model | Objective | Pixel Size (µm) (x*y*z) | Imaging Parameters | Channel | Wavelength (nm) | | Light Source | | Z-Stacks | | |
|---|---|---|---|---|---|---|---|---|---|---|---|---|---|
| | | | | | | | Excitation | Emission | Exposure time (ms) | Intensity (%) | Z-stack (µm) | Z-steps (µm) | Number of Slices |
| **Fig. 2, Sup. Fig. 1A/C/E** | **Hippocampus Cell Counts & Intensity** | Zeiss-Axio Imager.Z2 | Plan-Apocromat 20X/0.8 M27 | 0.173 µm x 0.173 µm | Tiled images with 30% overlap and stitched using Zen Blue. | **49 DAPI** | 353.0 | 465.0 | 40.0 | 25.0 | NA | NA | NA |
| | | | | | | **38 eGFP** | 488.0 | 509.0 | 390.0 | 32.0 | | | |
| | | | | | | **71 HcRed** | 590.0 | 618.0 | 40.0 | 25.0 | | | |
| | | | | | | **50 Cy5** | 650.0 | 673.0 | 225.0 | 25.0 | | | |
| **Fig. 3A-D, Sup. Fig. 1B/D/F** | **Dorsal Striatum Cell Counts** | Zeiss-Axio Imager.Z2 | Plan-Apocromat 20X/0.8 M27 | 0.173 µm x 0.173 µm | Tiled images with 30% overlap and stitched using Zen Blue. | **49 DAPI** | 353.0 | 465.0 | 40.0 | 25.0 | NA | NA | NA |
| | | | | | | **38 eGFP** | 488.0 | 509.0 | 390.0 | 32.0 | | | |
| | | | | | | **71 HcRed** | 590.0 | 618.0 | 40.0 | 25.0 | | | |
| | | | | | | **50 Cy5** | 650.0 | 673.0 | 225.0 | 25.0 | | | |
| **Fig. 3E** | **Colocalization Analysis** | Zeiss-Axio Imager.Z2 | Plan-Apocromat 40X/0.95 Korr M27 | 0.086 µm x 0.086 µm x 0.340 µm | Subtract background = 20 pixels, Average projection, analyzed with JaCoP on FIJI. | **49 DAPI** | 353.0 | 465.0 | 40.0 | 25.0 | **14.96** | **0.34** | **45** |
| | | | | | | **38 eGFP** | 488.0 | 509.0 | 275.0 | 32.0 | | | |
| | | | | | | **71 HcRed** | 590.0 | 618.0 | 40.0 | 25.0 | | | |
| | | | | | | **50 Cy5** | 650.0 | 673.0 | 225.0 | 25.0 | | | |
| **Fig. 4** | **Regression Analysis** | Zeiss-Axio Imager.Z2 | Plan-Apocromat 20X/0.8 M27 | 0.173 µm x 0.173 µm | Regions of interests intensities normalized before plotting. | **49 DAPI** | 353.0 | 465.0 | 40.0 | 25.0 | NA | NA | NA |
| | | | | | | **38 eGFP** | 488.0 | 509.0 | 390.0 | 32.0 | | | |
| | | | | | | **71 HcRed** | 590.0 | 618.0 | 40.0 | 25.0 | | | |
| | | | | | | **50 Cy5** | 650.0 | 673.0 | 225.0 | 25.0 | | | |
| **Fig. 5B, Sup. Fig. 2A-C** | **Microglia Cell Counts & Intensity** | Zeiss-Axio Imager.Z2 | Plan-Apocromat 20X/0.8 M27 | 0.173 µm x 0.173 µm | Tiled images with 30% overlap and stitched using Zen Blue. | **49 DAPI** | 353.0 | 465.0 | 26.0 | 25.0 | NA | NA | NA |
| | | | | | | **50 Cy5** | 650.0 | 673.0 | 200.0 | 25.0 | | | |
| **Fig. 5C-K** | **Microglia Morphology** | Leica LSM 880, AxioObserver | Plan-Apochromat 63x/1.40 Oil DIC M27 | 0.22 µm x 0.22 µm x 0.30 µm | Subtract background = 50 pixels, Average 2D projections. | **49 DAPI** | 405.0 | 455.0 | 1.02 µs/pixel | 2.2 | **15.00** | **0.30** | **51** |
| | | | | | | **38 eGFP** | 488.0 | 537.0 | 1.02 us/pixel | 2.0 | | | |
